# Fluctuations in arousal reflect latent state transitions that facilitate behavioral optimization

**DOI:** 10.1101/2025.02.06.636887

**Authors:** Tiantian Li, Harrison Marble, Timothy Chen, Niloufar Razmi, Matthew R Nassar

## Abstract

People are often faced with surprising events that defy expectations. Such events elicit transient activity in the locus coeruleus/norepinephrine system and elevation of peripheral arousal markers including pupil dilation and the EEG P300, but the function of these signals remains unclear. We propose that they reflect latent state transitions that dynamically control the mental context governing learning and perception. We tested and confirmed five preregistered predictions of this theory using EEG and pupil measurements collected from people performing a color prediction and reproduction task. Task latent state transitions elicited pupil dilation and amplified event-related potentials including the P300. These EEG and pupil measures related to behavioral signatures of latent state updating, including reduced bias and bidirectional adjustment of learning, both across trials and individuals. Our findings support the theory that LC/NE-linked arousal systems optimize behavior by signaling environmental transitions to facilitate rapid adjustments of mental context.

## Introduction

Learning in a complex and dynamic world is difficult and inevitably involves violations of our expectations. For example, when visiting a familiar restaurant, we enter with an expectation about the quality of food, but occasionally these expectations will be challenged by discordant perceptual information (e.g. a spicier than expected meal). Such surprising events are accompanied by transient arousal responses evident in peripheral markers such as pupil dilation (Alamia et al., 2019, Nassar et al., 2012, Preuschoff et al., 2011, Zhao et al., 2019). While previous work has made great strides in uncovering the biological bases of transient arousal responses (Devauges & Sara, 1991; Ghosh et al., 2021; Lee et al., 2018; Nassar, 2024; Nassar et al., 2012; Nieuwenhuis, 2011; Nieuwenhuis et al., 2005; Vazey et al., 2018) their function and contributions to behavior remain issues of ongoing scientific debate.

Transient fluctuations in arousal are mediated in part by the locus coeruleus/ norepinephrine (LC/NE) system. The LC/NE system in humans is composed of fewer than one hundred thousand NE-releasing neurons with cell bodies located in the LC that project broadly, providing the majority of NE modulation to the brain (Aston-Jones & Cohen, 2005; Nieuwenhuis et al., 2005; Sara, 2009). Transient release of NE leads to systematic changes in cortical processing (Bouret & Sara, 2005; Pineda et al., 1989; Vinck et al., 2015), large scale electrophysiological signatures reminiscent of the P300 orienting response (Nieuwenhuis, 2011; Nieuwenhuis et al., 2005; Vazey et al., 2018), and peripheral markers of arousal such as pupil dilation (Joshi et al., 2016; Joshi & Gold, 2021; Lloyd et al., 2023; Reimer et al., 2014). These changes in arousal are accompanied by altered behavioral performance in domains such as learning and perception (de Gee et al., 2017, 2020; Devauges & Sara, 1990; Ghosh et al., 2021; Krishnamurthy et al., 2017; Nassar et al., 2012; Nieuwenhuis, 2011; Urai et al., 2017; Yu & Dayan, 2005). Despite the frequency with which such fluctuations in arousal occur during everyday life, and the consequences that these fluctuations have on neural processing and behavior, the exact function that they serve remains unknown.

Recently we proposed a theory for how arousal systems are modulated to optimize behavior across domains (Li et al., 2023; Nassar, 2024; Razmi & Nassar, 2022). Our theory posits that transient activation of the LC/NE system signals the need to update active representations of mental context, which allows the brain to avoid interference that would be caused by combining information across distinct contexts. This theory predicts that fluctuations in LC/NE, and their peripheral arousal signatures, should relate to optimization of both learning (Dayan & Yu, 2006; Jepma et al., 2016, 2018; Nassar et al., 2019; Nieuwenhuis, 2011; Yu & Dayan, 2005) and perception (de Gee et al., 2017; Krishnamurthy et al., 2017; Urai et al., 2017) in complex environments, as we describe in more detail below. Our framework builds on theoretical and empirical research suggesting that both learning and perceptual inference are improved through maintenance of explicit representations of the latent causal process giving rise to observable data, which we refer to here as latent states (Collins & Frank, 2013, 2016; Gershman & Niv, 2013). In complex environments, effective learning and perception requires updating such latent states in real-time, in order to effectively attribute learning to the most relevant context, and to regularize perception in a context-appropriate manner (Collins & Frank, 2016; Nassar et al., 2010, 2019). In the restaurant example, if we discover that the normal chef is “out sick” (i.e. different latent state), it might affect how we perceive the food quality (i.e. perception) and it also might affect the degree to which we adjust our future expectations about the restaurant according to our current experience (i.e. learning). The timing of such latent state transitions is critical for both optimal learning and perception. If outdated latent states linger, then perception will be biased toward irrelevant expectations (e.g. when we entered the restaurant thinking we have the normal chef, our gustatory experience will be biased toward the one we expected – rather than accurately reflecting the food in front of us) and credit will be incorrectly assigned for learning (e.g. the wrong chef will be blamed for a bad meal). The need for latent state updating is often signaled by an external event that is surprising when viewed through the lens of the active latent state representation (e.g. the normal chef makes delicious soup, but today’s soup tastes terrible).

The locus coeruleus/NE system is well positioned to signal the need for latent state updating. The LC projects broadly throughout the brain and responds transiently to surprising events (Aston-Jones et al., 1999; Sara, 2009). While the effects of such neuromodulation are varied across tasks and systems, a confluence of data suggest that it may facilitate a rapid reorganization of cortical firing patterns, which has been referred to as a “network reset” (Bouret & Sara, 2005). We postulate that the functional role of this network reset is to facilitate a transition to a new active latent state (e.g. realizing this food was not made by the regular chef), thereby promoting an immediate disengagement from expectations stored in the associations linked to the previously active state (Li et al., 2023; Nassar, 2024; Razmi & Nassar, 2022). In doing so, such a mechanism would reduce biases to perceive new data in accordance with recent expectations (thereby optimizing perception) and ensure that credit for the change (e.g. bad meal) is attributed to the correct source (e.g. responsible chef), thereby optimizing learning.

Here we test five pre-registered predictions of the latent state theory of arousal by recording EEG and pupil diameter as proxies for LC-linked arousal systems during performance of a novel color prediction and reproduction task facilitating measurements of both bias and learning (Li et al., 2023). Our results confirm all five predictions, in particular demonstrating that both EEG- and pupil-derived arousal markers locked to stimulus presentation reflect latent state transitions (1), co-occur with reduced perceptual biases (2), and relate to learning in a context-appropriate bidirectional manner (3). Furthermore, pupil dilations aligned to predictions were consistent with reflecting an internally generated state transition signal necessary to avoid interference from oddball events (4). Finally, consistent with our theory of a common mechanism for the relationships of arousal to learning and bias, we show that individuals with stronger arousal modulation of bias also show stronger arousal modulation of learning (5). Taken together, our results provide strong support for the hypothesis that arousal systems optimize behavior by signaling the need to update active representations reflecting latent states of the world.

## Results

### A combined prediction and reproduction task for testing latent states theory of arousal

To test whether arousal systems relate to optimization of both learning and perception in line with our latent states theory, we developed a prediction/reproduction task that allowed us to measure both learning and perceptual bias in two different environments differing in their latent state dynamics. On each trial, participants were shown two colored patches and asked to reproduce the colors previously shown and then asked to predict the ones that would be shown next (Figure 1A). Participants performed two blocks of the task, each with its own generative environment that imposed a sequential structure that enabled participants to predict upcoming colors with some degree of reliability (Figure 1B). In the changepoint environment, colors were generated around a mean that changed occasionally (Figure 1B, top; see also Figure S5A). In the oddball environment, the mean drifted slightly on each trial, but colors were occasionally sampled uniformly from the entire color wheel, rather than from a distribution centered on the current mean (Figure 1B, bottom; see also Figure S5B). Pupil diameter and EEG data were recorded continuously as participants performed the task, allowing us to measure fluctuations in arousal, and serving as proxies for underlying LC/NE activation. A key advantage of this task is that it allowed us to measure behavioral constructs of interest on each trial (learning and perceptual bias), including those with latent state transitions (Changepoints/Oddballs; Figure 1C), allowing us to test if and how behavior changes with transient signatures of arousal (measured through pupil diameter and EEG).

**Figure 1:**
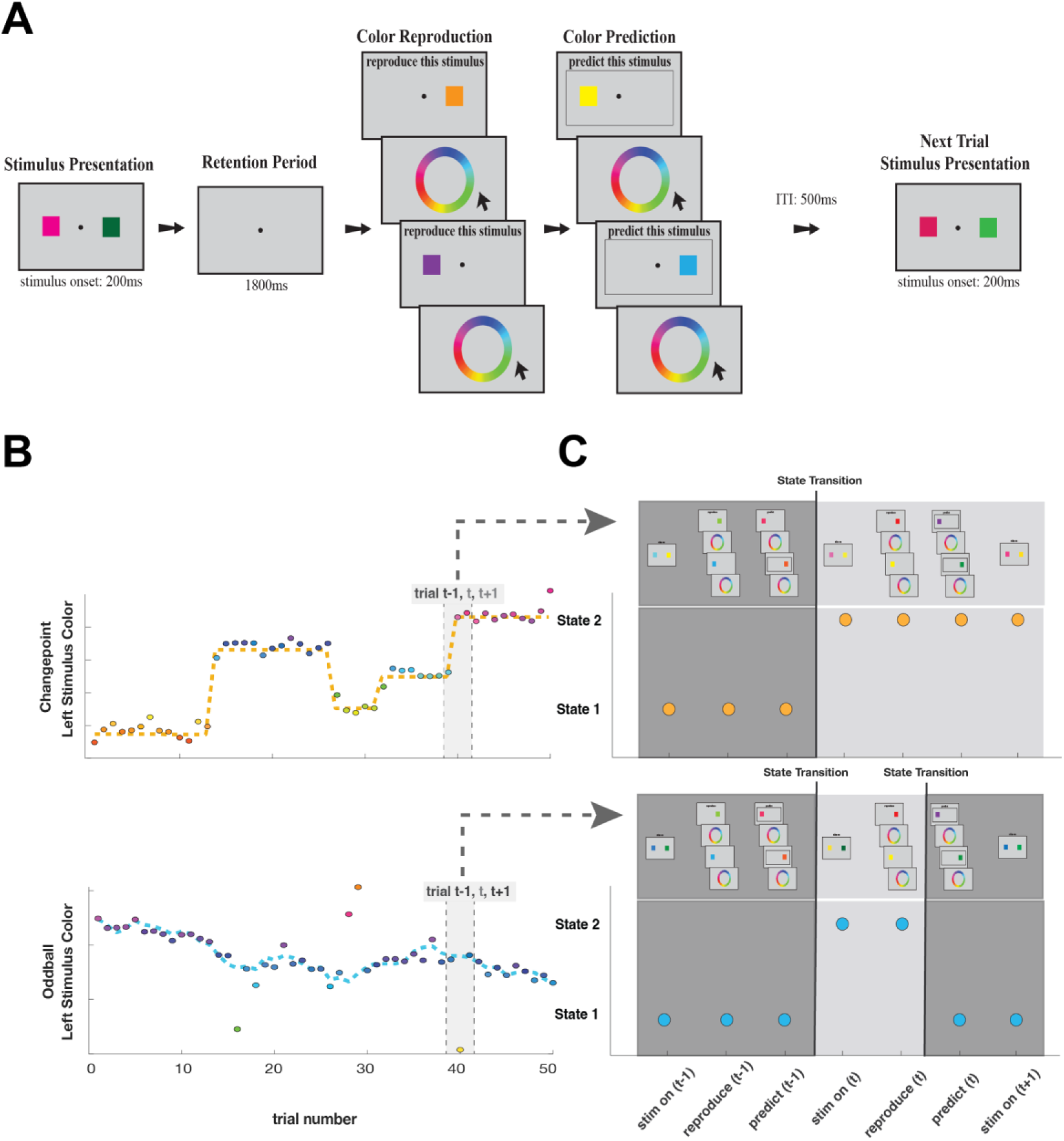
Task schematics. **(A)** On each trial, participants go through three main phases: stimulus presentation, color reproduction, and color prediction. **(B)** Examples of how the colors of one stimulus evolve across trials in the two different environments. As shown by the colors of circles in this figure, stimuli with close values in degree (y-axis) are similar in color. **(C)** A detailed look into latent state transition structures across two conditions at each task step before and after a surprising trial. For example, in B, both changepoint and oddball conditions have trial t as a surprising trial. In the changepoint environment, latent state transitions at stimulus onset on trial t and stays in the newly transitioned state until the next surprise stimulus. In the oddball condition, after transitioning to a new state with surprising stimulus, the state transitions back to its previous state after color reproduction in trial t to prepare for the next stimulus at trial t+1. So, the key difference between the two conditions is that changepoints give rise to persistent changes in latent state, whereas oddballs lead to a transient change in latent state and rapid reversion to the previously active latent state.

### Participants combined priors with sensory information to improve reproductions

In theory, combining prior expectations with internal representations of sensory information can allow for more accurate perceptions, particularly in cases where such priors are reliable predictors of future stimuli (Knill & Pouget, 2004; Knill & Richards, 1996). In particular, Bayesian inference would stipulate perceptions to reflect a weighted average of prior expectations and internal representations of sensory information:

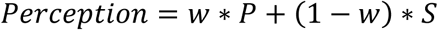

Where P is prior expectation, S is the sensory information, and the weights (w and 1-w) are proportional to the relative precision associated with each source of information. Note that this implies that perceptions should be biased towards expectations to the degree, w, to which those expectations are informative about the observed stimulus. To assess whether participants were able to capitalize on prior expectations to improve their task performance, we first compared reproduction errors observed in our primary task to those from an unstructured color reproduction control block, in which colors were uniformly sampled and thus prior information was uninformative. We found that reproduction errors were lower in our task than in the unstructured control block (Figure S1A; t = 4.74; dof = 62; p = 6.45*10^− 6^), suggesting that participants were able to capitalize on predictability in the task to improve the accuracy of their reproductions. Reproduction errors were also much lower than the prediction errors in the structured blocks (Figure S1B; t = 24.3; dof = 62; p = 2.26*10^−33^), suggesting that participants used sensory information to at least some degree. On average across trials, reproduction errors were similar to those expected from a Bayesian weighted combination of prior information (with reliability determined according to prediction errors) and sensory likelihood information (with reliability determined according to reproduction errors in the unstructured control block) supporting the idea that participants combined prior and sensory information in a near-optimal manner (Figure S1C; t = -0.95; dof = 62; p = 0.3477). Thus, on average across trials, participants combined prior and sensory information to achieve improved reproductions, allowing us to examine how the dynamics of task structure and arousal systems affect this process.

### (H1) Pupil and EEG signals reflect state transition probability

While prior expectations could be used to reliably predict upcoming colors most of the time, this was not true when the color sequence underwent a changepoint or oddball event, which we collectively refer to as state transitions. Our motivating theory posits that arousal systems must be engaged at such state transitions in order to prevent interference. Stemming from this theory, our first pre-registered prediction was that transient markers of arousal, including pupil diameter and the P300, should be increased in response to surprising stimuli that reflect likely state transitions in both task environments (Li et al., 2023).

We found evidence for such a relationship relating pupil dilations to likely state transitions. For each stimulus, we computed the probability that the stimulus reflected a state transition (state transition probability; STP) using a Bayesian ideal observer model calibrated to the appropriate task environment (oddball or changepoint). We then performed a running linear regression on each subject’s pupil data, using the average state transition probability of the two stimuli as a key explanatory variable. We found that pupils dilated substantially after stimulus presentation (Figure 2B), and that this dilation was larger for trials with higher STP. This enhanced pupil dilation occurred during a window 1.5-4 seconds after stimulus onset (Figure 2C; green). Consistent with previous work (Nassar et al., 2012; Krishnamurthy et al., 2017), we also found that baseline pupil size correlated with trial-to-trial entropy, which quantified uncertainty on the belief distribution over possible generative means prior to observing the stimuli (Figure 2A). Beyond this, we also found that entropy had a small effect on the magnitude of pupil dilations immediately after stimulus onset (Figure 2C; yellow). Thus, pupillometry data support the overarching prediction that arousal measures are elevated at state transitions and the uncertainty resulting from them.

**Figure 2:**
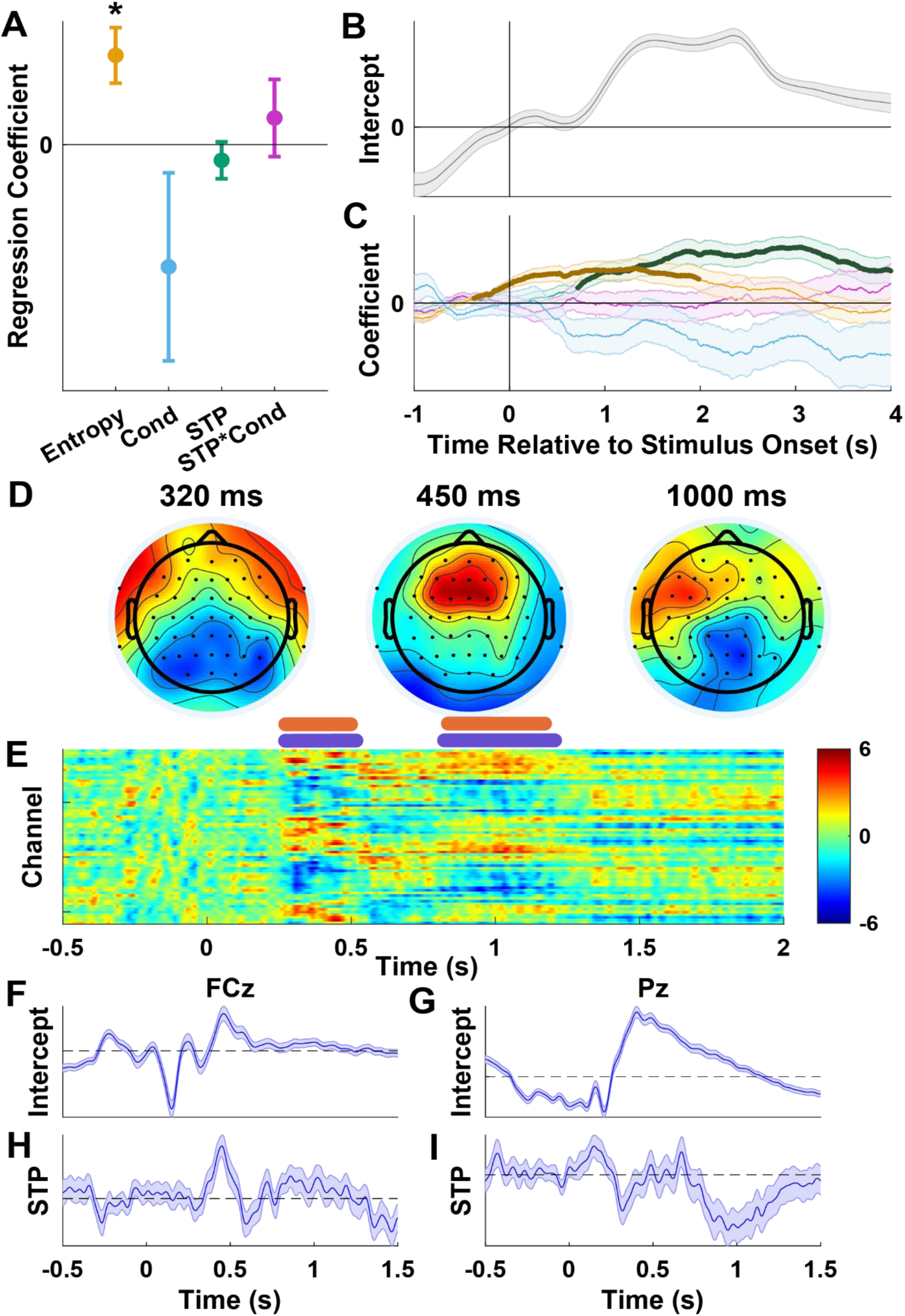
Pupil and EEG correlates of state transition probability (STP). **(A)** Points/lines reflect Mean/SEM regression coefficients reflecting relationship between task variables on baseline pupil diameter. Asterisks reflect significant t-test; Entropy t= 3.0, dof = 59, p = 0.0038. **(B)** Mean/SEM baseline subtracted pupil diameter plotted as a function of time after stimulus onset. **(C)** Lines/shading reflect mean/SEM regression coefficients reflecting relationship between task variables and evoked pupil response are plotted as a function of time after stimulus onset for STP (green), entropy (yellow), condition (blue) and STP*condition (purple). Dark points reflect significant temporal clusters after permutation tests controlling for multiple comparisons (STP cluster mass = 12673, p = 0.0004; Entropy cluster mass = 7982, p = 0.0048). **(D)** Scalp topography of t-statistics summarizing STP coefficients from mass univariate EEG regression is displayed for 320, 450 and 1000 ms post stimulus presentation. **(E)** Heat map displaying STP t-statistics from the same EEG regression, with channels reflected on the ordinate and time relative to stimulus onset reflected on abscissa. Horizontal lines indicate cluster timepoints surviving permutation testing for multiple comparison correction. Orange lines indicate positive clusters, blue lines indicate negative clusters. (Cluster Masses = 5349, 5979, 5449, 7647; p = 0.016, 0.0104, 0.0154, 0.0046) **(F-I)** Lines/shading reflect Mean/SEM coefficients for the intercept **(F/G)** and STP **(H/I)** terms of the EEG regression for channels FCz **(F/H)** and Pz **(G/I)**.

We next examined whether event-locked P300 responses, which are also thought to reflect a transient arousal and activation of the LC/NE system, were enhanced in response to surprising colors indicative of state transitions. To do so, we performed a running linear regression on the EEG data that included STP as an explanatory variable. We found that, on average, the stimulus onset induced P300-like responses at both frontal and parietal electrodes (Figure 2F&G). To test whether the magnitude of these responses was increased at state transitions, we examined the STP coefficients from the regression model and found four spatiotemporally clustered event-related potentials (ERPs) related to STP that survived multiple comparisons correction: a positive frontally localized signal resembling a P3a about 450 ms after stimulus onset, a negative parietal signal around the same time, a second separate negative parietal signal peaking around 900 ms after stimulus onset, and a second separate positive frontal signal peaking around 900 ms after stimulus onset (Figure 2D-E). Thus, our EEG data support the overarching prediction that the P300 reflects STP, but at a finer grain show that this is only true for the frontal P3a (as opposed to the parietal P3b, the later subcomponent of the P300), and also highlight three additional event-locked signals that reflect STP.

### Subjects reduce bias when presented with surprising stimuli

In stationary environments, people bias perception toward prior expectations, allowing them to achieve more accurate perceptions through Bayesian inference(Knill & Pouget, 2004; Knill & Richards, 1996). As described above, participants in our study appeared to combine prior knowledge and observed stimuli in order to improve performance on aggregate across all trials. However, priors become less useful at state transitions, when observed stimuli diverge from expectations. Optimal inference in complex environments requires decreasing bias in response to surprising stimuli if those stimuli are indicative of a change in the underlying state of the environment (Krishnamurthy et al., 2017). Thus, a signal that reflects state transitions should reduce reliance on priors, which could be measured as a reduction of bias toward them.

To test this idea, we first verified that participants in our study reduced their bias appropriately in response to surprising stimuli. Bias in our task can be thought of as the degree to which errors in reproduction match the errors made in prediction (oPE; Figure 3A). Consistent with Bayesian inference in dynamic environments, reproduction errors increased linearly with oPE over a wide range of moderate prediction errors, but the slope of this relationship decreased dramatically for extreme prediction errors, indicative of a reduction in bias for the most surprising outcomes (Figure 3B). This effect was present and comparable in both blocks and was even more pronounced for participants who made the smallest reproduction errors (Figure 3C). To quantify these effects across subjects we performed a linear regression that predicted reproduction error based on oPE and its interaction with STP. We found that prediction error had a positive effect on reproduction error, suggesting that participants were biased overall, but also that prediction error times STP had a negative effect on reproduction error (Figure 3D; t = -11.2; dof = 62; p =1.54*10^− 16^). Thus, participants biased their reproductions toward their predictions and reduced this bias in response to surprising stimuli reflecting likely state transitions.

**Figure 3:**
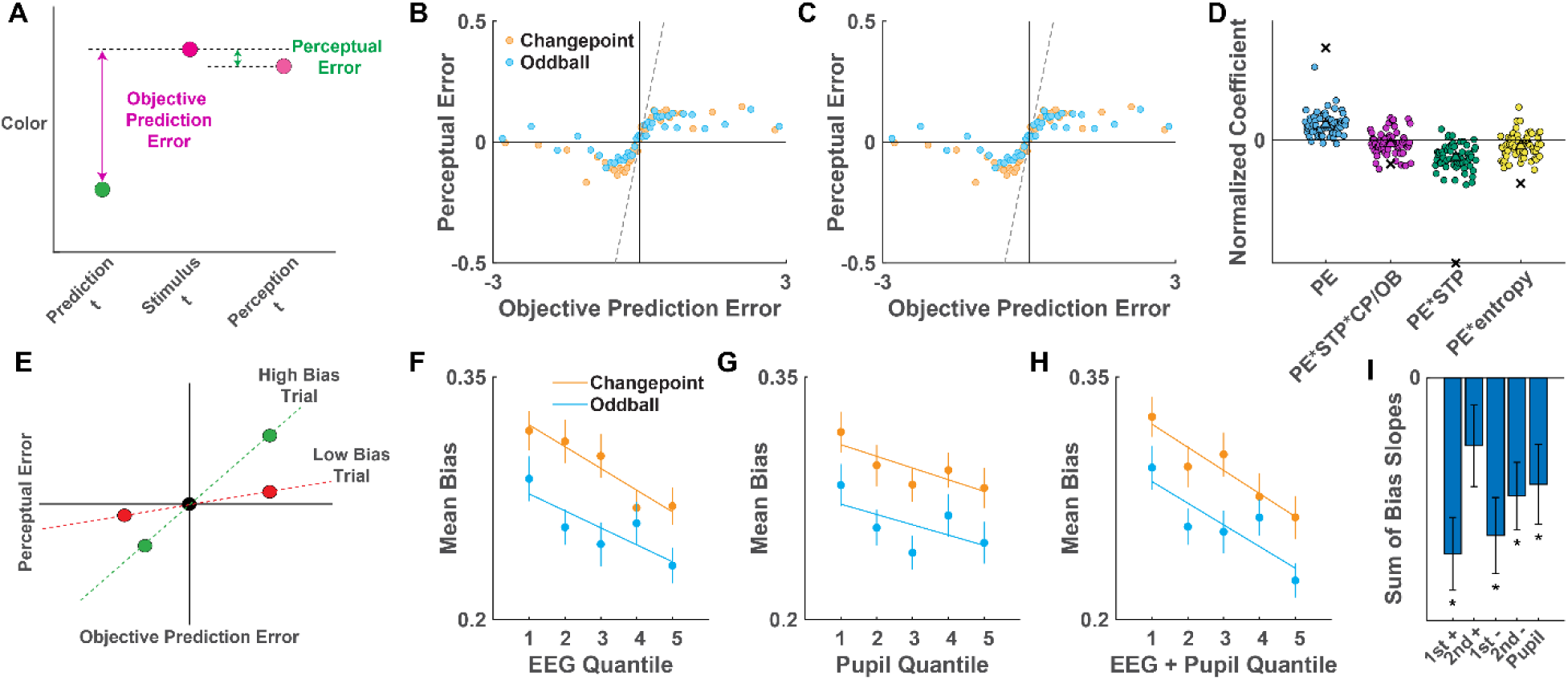
EEG and pupil signals that reflect state transitions correspond to reductions in perceptual bias. **(A)** Bias is calculated by dividing perceptual error by objective prediction error. **(B-C)** Mean perceptual error (ordinate) is plotted for bins of objective prediction error (abscissa) separately for changepoint (orange) and oddball (blue) conditions across all participants **(B)** and in just those participants achieving lower than median perceptual errors **(C).** Note that the positive relationship between prediction errors and perceptual errors is reduced for extreme errors, an effect that is even more pronounced in the highest performing participants. **(D)** Subject coefficients from a circular regression of trial-to-trial perceptual errors (Abscissa) are plotted separately for terms in the regression modeling capturing a fixed influence of prediction errors on perceptual errors (PE), as well as for interaction terms corresponding to dynamic adjustments of bias in according to STP (green), STP*Condition interaction (purple), or entropy (yellow). PE coefficients were positive across participants, reflecting a perceptual bias toward predicted colors (t = 10.4; dof = 62; p = 2.95*10^−15^), whereas PE*STP were negative, reflecting a reduction in perceptual bias for stimuli that reflected likely state transitions (t = -11.5; dof = 62; p = 5.62*10^−17^). **(E)** Schematic depicting how a single trial measure of bias was calculated for two stimuli presented on a given trial. A line is fit to the origin and both trial’s points on an objective prediction error/perceptual error plane, with its slope providing a single trial measure of bias. **(F-H)** increased strength of STP signals extracted from **(F)** EEG (Changepoint t = -3.90; dof = 56; p = 2.60 *10^−4^. Oddball t = -3.71; dof = 56; p =4.67 *10^−4^), **(G)** pupil (Changepoint t = -2.19; dof = 59; p = 0.0328. Oddball t = - 2.06; dof = 59; p = 0.0431), and **(H)** both (Changepoint t = -3.26; dof = 55; p = 0.0019. Oddball t = -4.06; dof = 55; p = 1.56 *10^−4^) corresponded to decreased bias. **(I)** EEG clusters and pupil signal are differentially related to bias, with the first negative ERP having the strongest relationship (t = -4.84, -1.66, -4.14, -3.51; dof = 55; p = 1.10*10^−5^, 0.1032, 1.20*10^−4^, 8.95*10^−4^)

### (H2) EEG and pupil signals that reflect state transitions are accompanied by reduced perceptual bias

Our ability to measure trial-to-trial fluctuations in perceptual bias allowed us to test a second preregistered prediction of our latent state theory of arousal: that arousal measures, if they signal the need to update latent states, should also be accompanied by reduced levels of perceptual bias resulting from that latent state update (Li et al., 2023). To test whether EEG- and pupil-derived state transition signals related to perceptual bias, we looked at how state transition signals on individual trials related to average bias on those trials. As stated above, we found that EEG and pupil signals related to STP (Figure 2C-E). To obtain single trial measures of these signals, we computed the dot product of the signal (EEG/pupil) recorded on a given trial with a template state transition signal (in time for pupil signal, in channels and time for EEG signal) derived from the thresholded t-map resulting from the STP regression (see methods). Thus, higher/lower values of this measure on a given trial reflected more/less STP-like arousal signaling. Since each trial had one EEG/pupil signal and two measures of bias, we combined the bias measurements into a single value by taking the slope of a line fit to a point on the origin and the two stimuli plotted on a coordinate system containing oPE as the abscissa and the reproduction error as the ordinate (Figure 3E).

Consistent with our hypothesis that P300 magnitude would correspond to reduced perceptual bias, we found that trials more closely resembling the EEG template for state transitions were accompanied by decreased bias in both the changepoint and oddball blocks (Figure 3F, Changepoint t = -3.90; dof = 56; p = 2.60 *10^−4^. Oddball t = -3.71; dof = 56; p =4.67 *10^−4^). However, this signal’s relationship to decreased bias was not solely driven by the P3a signal as we predicted, but rather related to each of the EEG-derived STP clusters (Figure 3I).

Pupil dilations were also related to reductions in bias. We regressed the trial-by-trial pupil-derived state transition signal against trial average bias and found that the physiological state transition signal was related to decreased average bias across both conditions (Figure 3G, Changepoint t = -2.19; dof = 59; p = 0.0328. Oddball t = -2.06; dof = 59; p = 0.0431). Thus, in both of our proxies for LC activity, we found that there is a relationship between state transition-mediated arousal and decreased bias. Combining across pupil and EEG measures revealed even more impressive relationships (Figure 3H, Changepoint t = -3.26; dof = 55; p = 0.0019. Oddball t = -4.06; dof = 55; p = 1.56 *10^−4^) that held up even after controlling for model-based measures of STP (Figure S2A; t = -2.38; dof = 55; p = 0.0207). When we investigated the relationship between bias and each individual cluster in the EEG state transition signal, we found that the frontal P3a, and both negative parietal signals were related to a decrease in bias across both blocks (Figure 3I). Thus, the arousal measures appear to provide subjective measures of surprise linked to state transitions and their behavioral consequences in terms of bias.

### Participants adjust learning in a context-appropriate manner

Latent state transitions also require separating information in the service of learning. A key advantage of our task is that by comparing predictions before and after observation of a stimulus, we can also determine how much a participant updated their predictions, which provides us with a single trial behavioral index of learning. In particular, learning in this task can be thought of as the degree to which predictions are updated (prediction update; the difference between prediction on trial t-1 and the prediction on trial t) for a given subjective prediction error (sPE; the difference between the color predicted on trial t-1 and the color perceived on trial t). Thus, the slope of the relationship between prediction updates and prediction errors provides a single trial measure of learning rate (Figure 4A; Nassar et al., 2012).

**Figure 4:**
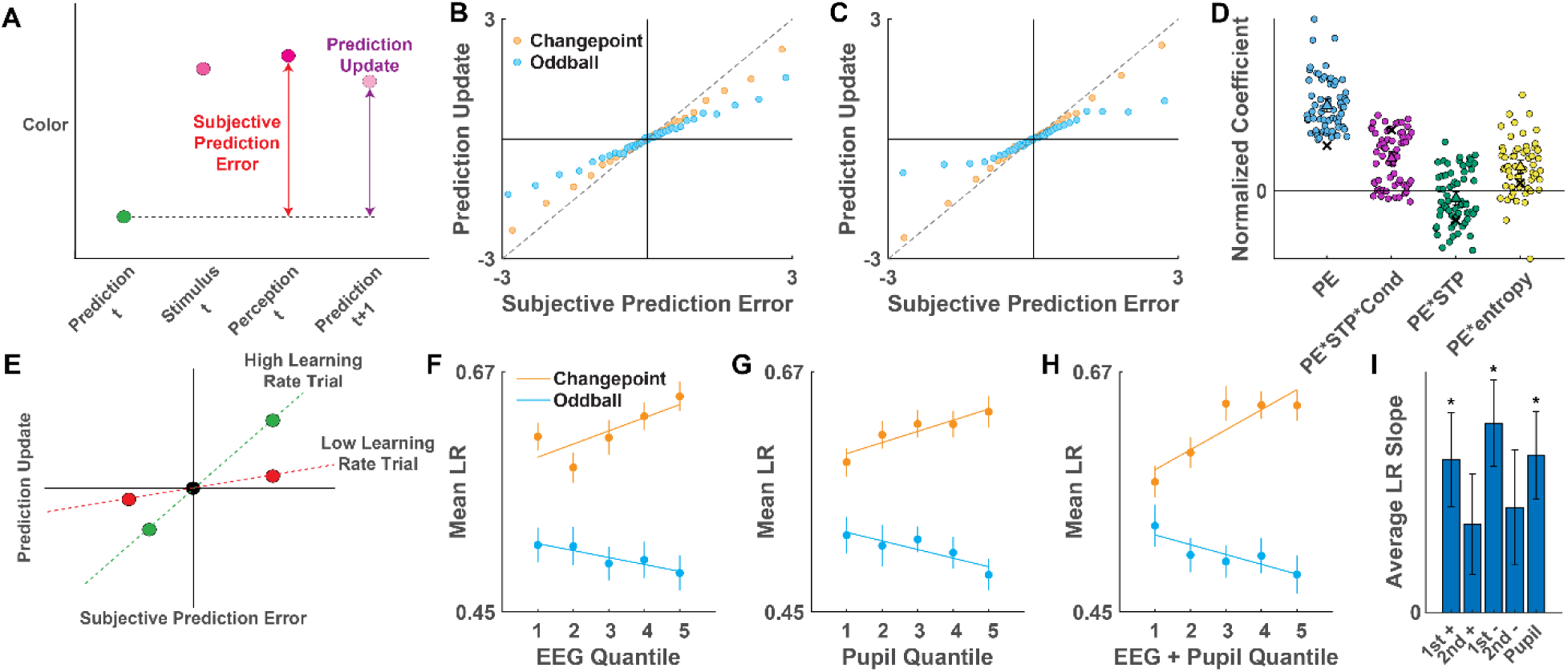
EEG and pupil signals that reflect state transitions are related to bidirectional adjustments in learning rate. **(A)** Learning rate is calculated as the degree of prediction update for a given subjective prediction error. **(B-C)** Mean prediction update (ordinate) is plotted for bins of subjective prediction error (abscissa) separately for changepoint (orange) and oddball (blue) conditions across all participants **(B)** and for just those participants who achieved lower than median average prediction errors **(C)**. Note that updates for extreme prediction errors diverge for the two conditions, with large prediction errors promoting increased learning in the changepoint condition and decreased learning in the oddball condition, an effect that is most obvious when looking at the subjects who made the most accurate predictions. **(D)** Subject coefficients from a circular regression of trial-to-trial prediction updates (Abscissa) are plotted separately for terms in the regression modeling capturing a fixed influence of prediction errors on updates (PE), as well as for interaction terms corresponding to dynamic adjustments of learning according to STP (green), STP*Condition interaction (purple), or entropy (yellow). PE coefficients were positive across participants (t = 25.2; dof = 62; p = 2.62*10^−34^), reflecting an overall tendency to update color predictions toward recently observed colors, as were PE*STP*Condition coefficients, reflecting the tendency of participants to increase learning in the face of changepoints but decrease it in the face of oddballs (t = 9.57; dof = 62; p = 7.91*10^−14^). (E) example calculation of trial mean learning rate (LR). A line is fit to points corresponding to the prediction update (ordinate) and prediction error (abscissa) for each of the two stimuli, as well as a third point corresponding to the origin. The slope of this line is taken to reflect the average learning rate for the trial. **(F-H)** increased strength of state transition signals from (F) EEG (Changepoint t = 3.08; dof = 56; p = 0.0038. Oddball t = -2.41; dof = 56; p = 0.0193), (G) pupil (Changepoint t = 3.06; dof = 59; p = 0.0033. Oddball t = -2.42; dof = 59; p = 0.0188), (H) or both measures combined (Changepoint t = 4.89; dof = 55; p = 9.24*10^−6^. Oddball t = -2.19; dof = 55; p = 0.0330) was conditionally related to single trial learning, increased state transition signaling reflecting more learning in the changepoint condition and less learning in the oddball condition. **(I)** EEG clusters and pupil dilation are differentially related to learning, with the first negative ERP having the strongest relationship (t = 3.25, 1.76, 5.04, 1.83; dof = 56; p = 0.0020, 0.0847, 5.30*10^−5^, 0.0723)

Consistent with normative prescriptions for learning (Nassar et al., 2019), participants updated predictions in response to sPE, and modulated their learning rate for large prediction errors differently across the two environmental contexts. For moderate prediction errors, participants increased their updates roughly in proportion to sPE in both changepoint and oddball conditions (Figure 4B). However, for extreme prediction errors, participants updated their predictions more in the changepoint than in the oddball block (Figure 4B). This effect was even more pronounced in the participants that made the most accurate predictions (Figure 4C). To quantify this effect for individual participants, we performed a linear regression that explained prediction update in terms of sPE and several interaction terms that modulated the effect of sPE (see methods). Consistent with the binned data (Figure 4B&C) we found that sPE coefficients were positive, capturing the overall positive relationship between sPE and updating (Figure 4D; blue). In addition, we found that the sPE * STP * condition coefficient was positive, suggesting that the slope of the relationship between updating and sPE was increased on changepoint trials and decreased on oddball trials (Figure 4D; magenta). In addition, there was also a small but statistically significant negative effect of sPE * STP on prediction update, suggesting that participants reduced their learning more on oddball trials than they increased it on changepoint trials (Figure 4D; green), as well as a positive sPE * entropy effect, suggesting that participants learned more during periods of uncertainty (Figure 4D; yellow). The Bayesian ideal observer model showed the same pattern of coefficients (Figure 4D; x markers) supporting the idea that participants adjusted their learning in a normative manner.

### (H3) Transient arousal signatures reflect bidirectional adjustments of learning

Our motivating theory proposes that arousal systems are transiently activated to signal the need for latent state transitions, and that such latent state transitions in turn produce contextual effects on learning rate; namely, increased learning from changepoints and decreased learning from oddballs (Razmi & Nassar, 2022). This leads to our third preregistered prediction: that arousal measures should relate contextually to learning behavior, corresponding to increased learning in response to changepoints, but decreased learning in response to oddballs (Li et al., 2023). To test this prediction, we extracted single trial measures of our arousal signatures as described above. We extracted a single trial measure of learning rate by combining learning rates corresponding to the two stimuli shown on a given trial into a single value corresponding to the slope of prediction updates relative to the sPE (Figure 4E). We then regressed our arousal measures (EEG/Pupil) onto trial average learning rates.

We found that both physiological markers for state transitions related contextually to learning behavior in the manner we predicted. Specifically, single trial ERPs that best matched the state transition template (including larger P300 responses) co-occurred with larger learning rates in the changepoint condition, but lower learning rates in the oddball condition (Figure 4F; Changepoint t = 3.08; dof = 56; p = 0.0038. Oddball t = -2.41; dof = 56; p = 0.0193). Similarly, the state transition-related pupil dilation was positively related to learning in the changepoint block and negatively related to learning in the oddball block (Figure 4G; Changepoint t = 3.06; dof = 59; p = 0.0033. Oddball t = -2.42; dof = 59; p = 0.0188). Combining the pupil- and EEG-derived state transition signals together yielded even stronger relationships to learning behavior (Figure 4H; Changepoint t = 4.89; dof = 55; p = 9.24*10^−6^. Oddball t = -2.19; dof = 55; p = 0.0330) that capitalized on similar relationships across all physiological STP signals (Figure 4I). The combined physiological signal was bidirectionally and contextually related to learning even after statistically controlling for objective STP (as computed by a Bayesian ideal observer model), suggesting that it provides a subjective measure of state transition probability that is closely linked to adjustments of learning behavior (Figure S2B; t = 3.11; dof = 55; p = 0.0030).

### (H4) Pupil dilations reflect an internally generated latent state transition

Our latent states theory of arousal makes different learning predictions for changepoint and oddball environments because these environments differ in their transition structure; namely, a changepoint reflects a deviation from expectations that is likely to persist for many trials, whereas an oddball reflects a deviation that is unlikely to. Or, through the lens of latent states, the key difference between oddballs and changepoints is that an additional latent state transition is required after an oddball event, but prior to making a prediction for the next trial (Figure 5A&B). This transition is adaptive for learning in that it allows the oddball to be treated differently from both the outcomes that preceded it, and also the outcomes that follow it. Since this transition is not locked to an external event, we anticipated that its timing would vary across trials and individuals, making it difficult to quantify with an event locked ERP (Li et al., 2023). However, pupil dilations, which provide a lagged and temporally smeared reflection of transient arousal responses, provide an ideal way to test this idea. In particular, they allow us to test our fourth pre-registered prediction, that pupil dilations between perceptual reports and the subsequent prediction will be greater for oddballs than for changepoints (Li et al., 2023).

**Figure 5:**
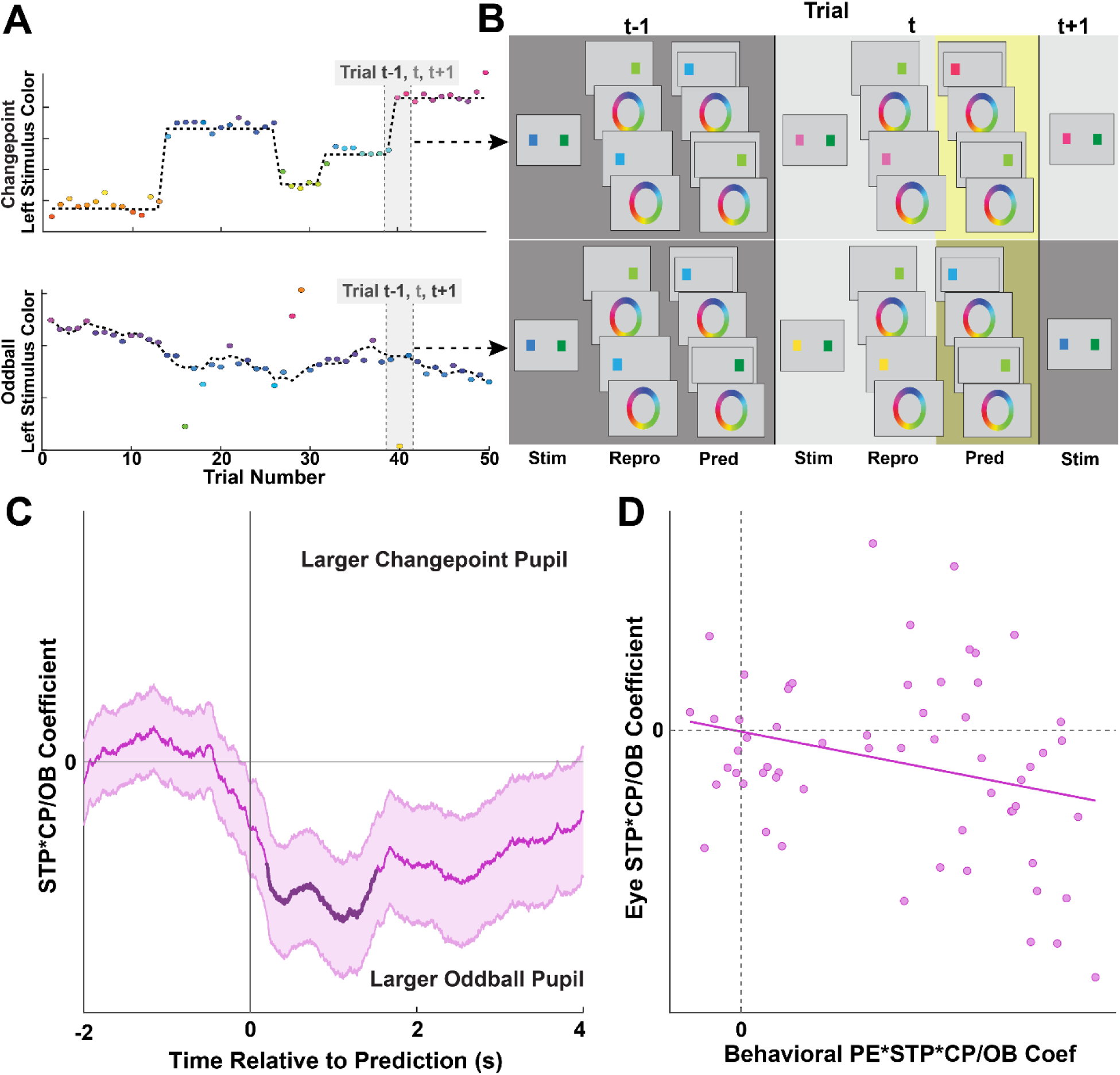
Delayed pupil dilations on oddball trials are consistent with reflecting an internally generated latent state transition. **(A)** Trial sequence containing a state transition in both changepoint (top) and oddball (bottom) conditions. **(B)** Task sequence with appropriate reproductions and predictions for the trials before during and after a state transition trial, showcasing that changepoints require one state transition while oddballs require two, with the second transition reflecting a return to the “normal” state (background color reflects state). **(C)** Mean/SEM STP*condition coefficients from a running linear regression of pupil size locked to the subsequent prediction phase of each trial. Dark points reflect significant clusters after permutation testing for multiple comparisons correction (STP*CP/OB cluster forming threshold = 0.025, Cluster mass = 3600, p = 0.0384). Decreased coefficients at the time of the subsequent prediction suggest a second pupil dilation occurring during the prediction phase after an oddball event. This supports the idea that pupil dilation is related to both internally generated and externally presented state transitions. **(D)** Correlation between participants’ behavioral PE*STP*CP/OB coefficient from behavioral regression and participants’ STP*CP/OB pupil regression coefficient during time window of significance. Those with stronger oddball effects in pupil also tended to have greater learning differences between changepoint and oddball trials (Pearson’s t-test: r = - 0.26; dof = 59; p = 0.0434).

We tested whether pupil dilations were greater for oddballs than changepoints when aligned to the subsequent prediction. Consistent with our hypothesis, there was a negative effect of STP * condition on pupil size, indicating that pupil dilations were larger following state transitions in the oddball condition (Figure 5C; Cluster forming threshold = 0.025, Cluster mass = 3600, p = 0.0384). This effect was strongest for the participants who showed the greatest modulation of learning behavior across the two conditions. There was a weak but significant correlation between this STP * condition coefficient in the pupil regression and the PE * STP * condition coefficient from our behavioral regression, showing that those with larger learning rates on changepoints and lower learning rates on oddballs also had larger pupil dilations after oddball events (Figure 5D; Pearson’s t-test: r = -0.26; dof = 59; p = 0.0434). Taken together, these results are in line with our prediction that individuals who understand the temporal structure of the two environments undergo a second internal latent state transition corresponding to the anticipation of a return to the previous state after an oddball event.

It is noteworthy that the timing of this effect occurred 200-1500 ms *after* prediction, raising questions about whether it might reflect an underlying neural process sufficiently early to influence that prediction (Figure 5C). Given that LC spiking precedes peak pupil dilation by 600-800 ms depending on experimental conditions in monkeys (Joshi et al., 2016), this pupil dilation would correspond to a difference in LC activity across our conditions emerging 400 to 600 ms before participants indicated their updated prediction, suggesting that, at least in theory, the effect we observe in the pupil could reflect firing in LC early enough to influence the prediction. Consistent with this idea, reanalyzing the same pupil date in terms of their derivative, which follows LC activation with a shorter lag, yielded effects that preceding, rather that following, the prediction (Figure S3). The oddball > changepoint effect was more tightly locked to the prediction than to the reproduction that preceded it, as the same analysis aligned to the preceding reproduction yielded no differences. Taken together, these results support a nuanced prediction of our overarching theory that arousal systems optimize behavior by signaling latent state transitions, in particular that oddball events elicit an additional delayed pupil response that would be consistent in timing with a second internally generated state transition.

### (H5) Individual differences support a single mechanism linking arousal to bias and learning

If arousal influences behavior by acting to signal or promote latent state transitions, then it should affect both bias and learning, and thus the degree to which it is engaged to modulate bias should be predictive of the degree to which it is engaged to modulate learning. Since subjects vary in the strength of their relationships between arousal and learning/bias, we could use these individual differences to test our theory that both arousal-behavior effects emerge as a result of a single unitary phenomenon. If learning and bias are both driven by the same arousal-mediated computation, then the degree to which arousal signals reflect learning in an individual should be related to the degree to which the same signals reflect perceptual bias in that individual. Taking the bidirectional learning relationships into account, this leads to our fifth and final preregistered prediction: there should be a stronger difference in the learning vs arousal slope in the changepoint and oddball blocks for individuals with a stronger relationship between bias/arousal (Figure 6A; Li et al., 2023).

**Figure 6:**
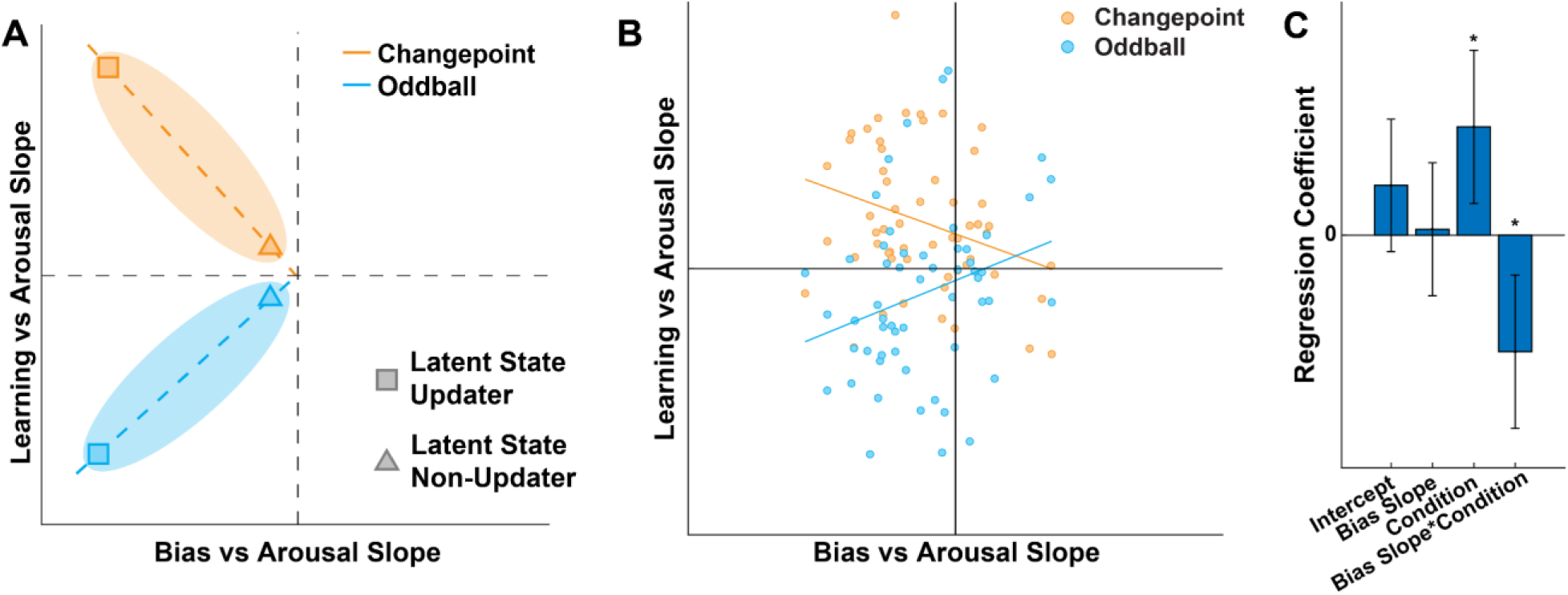
Individual differences between participants show that those with strong relationships between learning and arousal tend to have strong relationships between bias and arousal. If the arousal markers relate to both perceptual bias and learning because they reflect state transitions that influence both, then the degree to which a participant shows one relationship should predict the degree to which they show the other. **(A)** hypothesized relationships between learning vs arousal slope (ordinate) and bias vs arousal slope (abscissa) for changepoint (orange) and oddball (blue) blocks. **(B)** Participant data, blue points indicate subjects’ slopes in the oddball block, orange points indicate slopes in the changepoint block. While learning vs arousal slopes were individually calculated within each block, bias vs arousal slopes were calculated across both blocks. **(C)** Regression results against learning slopes. The significantly negative effect of bias slope * condition suggests the learning and bias signals share a similar origin, as subjects with stronger learning/arousal signals also have strong bias/arousal signals (Condition t = 2.79; dof = 108; p = 0.0062. Bias slope * condition t = -3.01; dof = 108; p = .0033)

Examining individual differences in arousal’s relationship to learning and bias provided support for these basic predictions. There was a negative relationship between learning/arousal relationship and bias/arousal relationship in the changepoint block, implying that in the changepoint block participants with a stronger negative relationship between bias and arousal also had a stronger positive relationship between learning and arousal (Figure 6B). To quantify the effect of bias/arousal slope on learning/arousal slope, we performed a regression on learning/arousal slope. We found that there was a positive effect of condition on learning/arousal slope, implying that, as predicted, the learning/arousal slope was higher for the changepoint than the oddball condition. Furthermore, the relationship between arousal and bias is differentially related to that between arousal and learning depending on condition, with the relationships in the changepoint condition more positive and those in the oddball more negative (Figure 6C; Bias slope * condition t = -3.01; dof = 108; p = .0033). This relationship was maintained even when STP was regressed out of the joint pupil-EEG arousal signal, trial average bias, and trial average learning rate, which suggests that these relationships are not solely due to the inherent relationships between STP, learning rate, and bias (Figure S2C-D; Bias slope * condition t = -2.67; dof = 108; p = .0087). Taken together, these results provide evidence for a common source of arousal’s effects on learning and bias, a key prediction of our latent state theory of arousal.

## Discussion

Our results provide support for the theory that arousal signals in the brain signal the need for updating of mental contexts that serve as internal latent states, leading to optimization of perception and learning. Pupil dilations and EEG signals including the P3a, a subcomponent of the P300, were elevated in response to surprising outcomes and reflected state transition probability (STP) across different environmental contexts. We found that pupil and EEG-derived STP signals were related to ongoing perceptual processing, with increased arousal signatures linked to lower bias in color reproductions. Furthermore, we found that the same transient arousal signatures were bidirectionally linked to learning, predicting heightened learning in an environment with changepoints, and reduced learning in an environment with oddballs. Furthermore, we found that pupil dilations preceding predictions in the task were increased for oddballs relative to changepoints, consistent with reflecting the internally generated latent state transition necessary to return to the latent state associated with typical trials before making a prediction. The size of this dilation event was also found to be related to a subject’s learning rate differences between oddballs and changepoints. Finally, we showed that individual differences in the relationship between arousal and perception, bidirectionally and contextually predict those between arousal and learning, consistent with both relationships being mediated by a single common cause, the degree to which participants relied on arousal systems to reset latent states at discontinuities in the data stream. Taken together, these results provide strong support for our latent state theory of arousal: that transient signatures of arousal signal the need to update internal latent state transitions, which in turn leads to optimization of both learning and perception to minimize the negative consequences of interference on behavior.

Our theory and results provide a unifying framework for two prominent lines of inquiry related to arousal and its effects on behavior. First, it accounts for relationships between arousal and learning, in particular findings that pupil dilations and P300 signals predict the degree to which people use new experience to update their beliefs about the world (Browning et al., 2015; Fischer & Ullsperger, 2013; Jepma et al., 2016, 2018; Nassar et al., 2012). However, our results demonstrate the need for reinterpretation of these original results since they reveal that the relationship between arousal signals and learning are both bidirectional and contextual. The signals co-occur with increased learning when latent state transitions tend to persist (changepoints) but decreased learning when such transitions are fleeting (oddballs). Second, our work relates to a line of research showing that arousal signals correspond to reductions in perceptual and decision biases (de Gee et al., 2017; Krishnamurthy et al., 2017; Urai et al., 2017). Our empirical results expand on these lines of work by demonstrating parallels across EEG and pupil diameter in their relationships to both learning and bias, and our latent states theory not only provides an explanation for these relationships, but makes unique predictions about contextual signaling differences and individual differences that were confirmed in our dataset. Thus, a primary contribution of our study was to unify two lines of behavioral research into a single set of behaviors that correspond to latent state transitions and in turn relate them to transient markers of arousal. While our study relied on peripheral markers of arousal, in particular EEG and pupillary correlates of latent state transitions, we believe that our results are suggestive regarding the underlying function of the LC/NE neuromodulatory system. Our motivating theory is based heavily on known physiology, pharmacology, and anatomy of the LC/NE system. In particular, that neurons in LC respond to surprising stimuli irrespective of modality and project broadly to cortex and subcortical regions to release norepinephrine (1). In target regions, NE amplifies feedforward processing (Devilbiss & Waterhouse, 2011; Rodenkirch et al., 2019) and weakens local recurrent connections (Cools & Arnsten, 2022; Hasselmo et al., 1997), providing a mechanism through which active latent state representations might be “reset” according to new inputs. Furthermore, activation/inactivation of the LC/NE system facilitates/inhibits behavioral flexibility (Donchin & Coles, 1988a; Jepma et al., 2016). Thus, the LC/NE system has response properties consistent with signaling latent state transitions, effects in target regions capable of promoting them, and effects on behavior consistent with serving that computational role. Pupil diameter and the P300 are thought to be proxies for LC/NE system activation (Joshi et al., 2016; Joshi & Gold, 2021; Murphy et al., 2011; Nieuwenhuis et al., 2005; Vazey et al., 2018) and while these proxies are indirect and nonspecific, by proxy our results provide further support for the idea that the LC/NE system affects behavior by promoting latent state transitions. In particular, our behavioral analyses provide an incredibly specific characterization of the computational role of these proxy measures, and this characterization supports the latent state transition theory. We suspect that the underlying computation is carried out by the LC/NE system, and we hope that future work will employ direct measurement and/or manipulation of the LC/NE system to test this idea.

It is worthy of note that the latent state transition theory of LC/NE function is quite closely related to two other prominent theories of LC/NE function. First, the notion that neural networks are “reset” in response to LC/NE signaling in order to facilitate rapid adaptation, or the “network reset hypothesis” was proposed by Bouret and Sara in 2005(Bouret & Sara, 2005). Second, the idea that LC/NE might be recruited in response to surprising stimuli that indicate a likely transition in the environment “unexpected uncertainty theory” was proposed by Yu and Dayan in 2005 (Yu & Dayan, 2005). The latent state transition theory, which we used to generate our predictions and to which our experimental results lend considerable support, can be thought of as a combination of these two theories, whereby unexpected events are detected and used to reset neural networks that represent the latent state of the world. While the latent states theory is not mutually exclusive from these other theories, it has advantages in terms of specificity and generality. In particular, we view the latent states theory as complementing “network reset hypothesis” by providing more specific predictions about how exactly neural networks should be updated to optimize behavior (Bouret & Sara, 2005), and complementing “unexpected uncertainty theory” by generalizing it to a broader range of generative environments beyond those for which it was originally mathematically derived (Yu & Dayan, 2005).

While our results largely fell in line with the latent states theory that motivated our predictions, there were several findings that were unanticipated. First, the spatiotemporal profile of the state transition signal we observed in the EEG data differed from that observed in another task that included both changepoint and oddball conditions (Nassar et al., 2019). Here we observed only one spatiotemporal cluster that resembled the P300. While previous work identified a state transition signal that included both early frontal (P3A) and later parietal (P3B) components, in the current work we found that only the former was related to state transitions. It is noteworthy, however, that although previous work linked both the P3A and P3B to state transition signals, only the early component of the P300 (P3A) was related to learning after controlling for other factors including state transition probability itself. Taken together with our current findings, this suggests that the P3A component may play a more central role in the neural mechanisms linking inferred state transitions to their behavioral consequences (as compared to the P3B). It is also worthy of note that our task was fundamentally different than the predictive inference task previously used to identify relationships between latent state transitions and the P300, since participants needed to use working memory to store colors on each trial, irrespective of the level of surprise or the need for updating latent states. The P3B might reflect working memory updating (Donchin & Coles, 1988b), which is consistent with the large stimulus locked P3B responses, irrespective of trial type, observed in our study. Thus, one possibility is that, in absence of an explicit working memory demand, working memory updating systems can be recruited to participate in the updating of latent states, leading to relationships between the P3B and latent state updating in predictive inference tasks, but not working memory tasks, where these systems are otherwise engaged. Nonetheless, it remains unclear why, in our data, the global parietal signal was reduced on state transition trials. The heterogeneity of EEG signals identified in our study also raises further questions, in particular why frontal positive and parietal negative signals related to latent state transitions emerged approximately one second after stimulus presentation. The functionality of these signals was mixed, as the parietal negative clusters were related to behavior while there was less evidence in this regard for the late frontal positive cluster. We hope that future studies will further elucidate the different roles of these EEG responses in coordinating adaptive behavior.

One additional discrepancy between our results and those we predicted is related to the timing of the pupil dilation that we predicted to occur in response to internal state transitions on oddball trials. We hypothesized that this second oddball signal would occur between the perceptual report and the prediction, as our theory stipulates that participants would need to switch latent states during this period in order to make accurate perceptual reports and subsequent predictions. Instead, we found that the pupil signal occurred slightly after participants made their predictions, which we interpreted as occurring due to the delay between the LC signaling of the state transition and the subsequent pupil dilation event, which in monkeys has been estimated at around 800 milliseconds (Joshi & Gold, 2021). Nonetheless, the timing of this event was later than anticipated, and certainly raises questions about whether the arousal systems are activated before or after internal responses to latent state transitions, a question that would be best posed to answer using more direct and time resolved measures of LC/NE function.

In summary, our results support the overarching idea that the brain optimizes learning and perception through dynamic transitions of internal latent state representations. Transient markers for arousal, including pupil diameter and the P3A reflect these transitions and relate to learning and bias behaviors accordingly. These relationships appear functional, since they persist after controlling for objective measures of state transition probability, and are stronger in participants who modulated behavior more. Taken together our results support a role for arousal systems in coordinating the dynamics of ongoing context representations that are used to fine tune perception and learning.

## Methods

### Task Design

To test our theory that arousal systems signal latent state transitions and their downstream behavioral consequences, we designed a color perception and prediction task that operates in two distinct statistical contexts. On each trial, participants were presented with two colored-target stimuli for 200ms and subsequently prompted to reproduce each color after a delay period. The task was divided into four blocks. The first block was purely for practice as participants reproduced the displayed stimuli and received feedback on the accuracy of their perceptual reports. The colors in this block were randomly generated. The second block was a longer iteration of the basic structure of the first block, however participants do not receive accuracy reports on their individual color estimations (although they did receive feedback at the end of the block). Given that colors displayed in the second block were chosen randomly from a uniform distribution, behavioral data from this block was used as a control condition to better understand whether participants were able to exploit the structure in later blocks in order to improve their color reproductions.

In the third and fourth blocks, participants performed the same task but modified in two important ways. First, participants were asked to predict the colors of the stimulus that would be presented at each location on the following trial. Predictions were possible because colors at each location were generated through an independent sequential generative process following either the changepoint (CP) or oddball (OB) state transition structure. The generative structure was instructed and explained to the participant. The third and fourth blocks were counterbalanced for order, so if a participant experienced the changepoint condition in block 3 they would experience the oddball condition in block 4, or vice versa. Predictions and estimations were considered correct, and participants are awarded one point if the error fell below 30 degrees. If the error was greater than 30 degrees, participants were penalized with a deduction of 1 point. At the end of the study, participants received one bonus dollar for every 30 points that they scored up to 20 dollars (600 points). Both CP and OB blocks contained 120 trials, and the task lasted between 1.5 to 2 hours in total.

Colors in our task were chosen from a circle in CIELAB color space (radius in A*, B* = 50, white point = [110, 100, 90]) along a plane with fixed luminance (L*=60). Standardized colors were shown by characterizing the experimental display using methods standard in color perception research (Cohen et al., 1967; Schloss et al., 2018; Zhang & Luck, 2008). The CIELAB color space was chosen for its approximate perceptual uniformity, which allows us to interpret angular errors in terms of fixed units of perceptual difference. The luminance of our color circle was fixed to minimize stimulus-related variability in the pupillary light reflex. All stimuli were displayed on a background with chromaticity equal to the white point with luminance similar to the luminance of the stimuli (L*=50). Colors are referred to by angular position in the task for convenience. All human subject procedures were approved by the Brown University IRB.

### Task Generative Model

Stimulus colors for each location were determined according to one of two generative models reflecting either the changepoint or oddball condition (Figure S5). In both cases, a generative mean (*M*) with a value between 0 and 360 is sampled to reflect the underlying mean of the color distribution (*M*_*t*_ ∼ *U*(*0*,*360*)). *M* serves as the mean of a von Mises distribution across a color spectrum from which a color value (*X*) is drawn 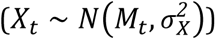, where 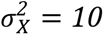. Beyond the observable aspects of the task, we assume that subjective internal representations of the stimulus value are corrupted by noise, and thus draw an internal representation (*Y*) from a normal distribution centered on *X*, which we refer to as sensory noise 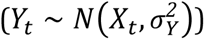. Thus, while X reflects the color shown in our task, Y reflects the subjective internal representation of that color accessible by the participant. In order to constrain the degree to which internal representations are corrupted, we set 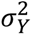 according to the measured variability in participant reports during the unstructured control task block. We found that the average estimation error on a trial was ∼31 degrees in the unstructured control block, and thus for our simulations we set *σ*^2^ to 31.

The generative mean (*M*) transitions across trials in one of two possible ways depending on the environment in each block. In the changepoint condition, a binary variable *S* is sampled according to a Bernoulli distribution with probability *H*, for hazard rate. If *S* is equal to 1, we draw the next generative from a uniform distribution. Otherwise, we draw the next generative mean from a normal distribution whose mean is the generative mean of the previous trial leading to the conditional sampling statement below:

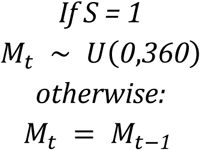

In either case, the true stimulus value is drawn from the current generative mean:

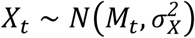

The generative model for the oddball condition differs from the changepoint condition in two important ways. First, the generative mean M transitions according to a Gaussian random walk across all trials 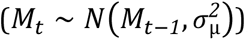, rather than only changing discrete changepoints. Second, the color on each trial is chosen conditionally according to S, such that if S is equal to one, then the color (true outcome: *X_t_*) is chosen uniformly across the entire color wheel, otherwise, the color will continue being drawn from the generative mean of the trial (*M_t_*). This gives rise to the following conditional sampling statements:

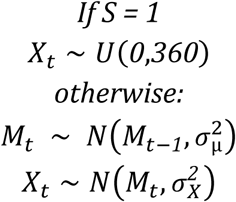

### Bayesian Inference Model

Our inference model assumes that participants understand the generative structure of the task, but only have access only to the observable parts of the generative model (*Y_1_, Y_2_, Y_3_… Y_t_*). We assume that subjects use this sequence of corrupted internal representations to infer the mean of the color distribution *M_t_*, which can be used as a prediction on each trial, and estimate the color of the observed stimulus *X_t_*, which can serve as a perceptual report on each trial.

The inference model can be arrived at by inverting the generative model (Figure S5) to infer *M* and *X* based on previous observations:

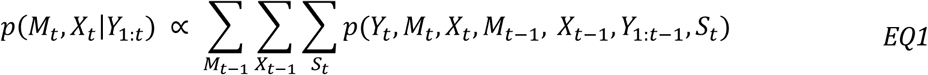

Note that this expression includes summations over the mean on previous trial (*M_t-1_*), the color on the previous trial (*X_t-1_*), as well as the possible state transitions on this trial (*S* = 1 corresponds to changepoint or oddball depending on condition, *S* = 0 corresponds to a standard transition). In the changepoint condition, conditional independencies from the generative graph can be exploited to yield the following factorization:

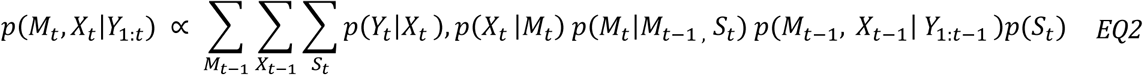

Where *S_t_* reflects the prior probability that any given trial will be a changepoint, referred to in the text as the hazard rate. Conditional independencies in the oddball condition, yield a slightly different factorization

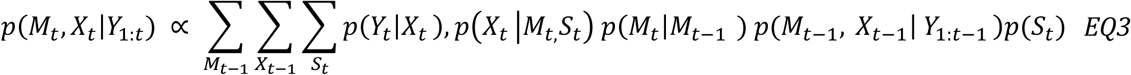

In both cases, the model infers *M_t_* and *X_t_* based on the observed history of *Y*, and this inference serves as a prior for a Bayesian update on the subsequent trial. The model produces an estimate of the stimulus shown by reporting the maximum posterior probability estimate of the stimulus (*X_t_*). Note that since inference includes not only the pure sensory observation *p*(*Y*_*t*_|*X*_*t*_) but also prior information based on learning of the underlying mean (terms 2-4 in Eq 3), the Bayesian inference model makes estimates about the stimulus that are biased toward its prior expectations, but to a degree that respects the generative structure of the task that includes occasional abnormal events (S). The inference model obtains its prediction of the next stimulus from the maximum posterior probability estimate of the generative mean (*M_t_*).

State transition probability estimates were computed as the integrated probability that a given observation reflects a latent state transition (*S_t_*), which reflects the model’s estimate of changepoint probability or oddball probability depending on the condition. This measure is monotonically related to standard measures of surprise, such as Shannon information, but differs in that it saturates at the point where a state transition is certain.

Trial-to-trial measures for state transition probability and learning extracted from the model were dissociable across environments: positively correlated in the changepoint environment while negatively correlated in the oddball environment. This feature allows us to leverage complementary environments that disentangle normative influences on learning and perceptual bias.

Previous studies have shown that uncertainty affects learning such that learning rate experiences slow decay after state transition. Here we measure uncertainty as the entropy on the model’s inference about the underlying mean, which is highest on trials after state transitions.

### Data Collection & Preprocessing

We collected behavioral, EEG (Brain Vision actiCHamp Plus 64, 1000Hz), and pupillometry (SR Research Eyelink 1000 Plus, 1000Hz) data from our subjects while they performed this task. In total, behavioral data was recorded from 63 subjects, 60 of which had usable pupil data, 57 of which had usable EEG data, and 56 of which had usable pupil and EEG data.

EEG preprocessing was performed in EEGLAB with the EYE-EEG extension for eye data-guided independent components analysis (ICA) (Dimigen 2020). We used a high pass filter of 0.1 Hz to remove low frequency noise, then segmented the data into 300 epochs each containing the 2 seconds before and after a trial. Each individual trial and channel data was then visually inspected and channels with persisting noise or artifacts were removed and interpolated. Trials containing excessive noise or artifacts were excluded. Subjects with >50% rejected trials or >25% rejected channels were excluded from further analysis. Next, we re-referenced each channel’s data according to the mean signal. Independent component analysis (ICA) was then performed on the data. We used the EYE-EEG extension to remove components with higher correlation to eye blinks and saccades than some threshold. We then low pass filtered the EEG data at 30 Hz. Cleaned data was then inspected manually to identify and remove any remaining trials with excessive noise or artifacts.

Eye-tracking data preprocessing was done using a custom MATLAB script. First, we removed any data from eyes that were poorly recorded. For subjects where one eye’s data was markedly worse than the other, only the data from the eye with better recordings was maintained. Next, we interpolated eye data during blinks and during a window 150 ms before and after each blink. This also filled in any gaps in eye data recording that were created by the eye tracker failing to track either eye. The data was then averaged across both eyes and epoched according to the time window that was being investigated. Trials with >20% missing data were rejected for all pupil alone analyses. Trials with >50% missing data were rejected for joint pupil-EEG analyses. We set this higher threshold for the joint analyses to retain as many trials as possible, since many trials had already been rejected during the EEG preprocessing pipeline. Subjects with more than 50% missing data on 50% or more of the trials were rejected from eye data analysis entirely.

### Behavioral Data Analysis

In order to examine how latent variables from our model influence learning and perceptual bias, we generated two linear regressions. Through these regressions, we analyzed how state transition probability modulates learning and perceptual bias.

To examine learning, we measured trial-to-trial prediction update computed as: color prediction(t+1) – color prediction(t). Learning rate was measured at the individual subject level as the linear relationship between prediction update and prediction error. We calculate this learning rate in two ways. The first is simply to divide prediction update by subjective prediction error. For this method, to account for the circular nature of the color space, trials with updates near 180 degrees were adjusted so the update and error had the same sign. After making this adjustment, we found that model learning rates correlated much more strongly with task variables such as STP. The second method of measuring learning rate is a linear regression model where we can measure effects of certain variables on update. Model learning rates were also calculated using this method. Prediction error (PE), in this case, was defined as the subjective prediction error (sPE), which is the difference between the trial stimulus color reproduction, a reflection of what the participant believes they saw, and the color prediction on the trial before stimulus onset. In order to understand how learning rate differs across different conditions, we performed the following regression on both model and subject behavior data:

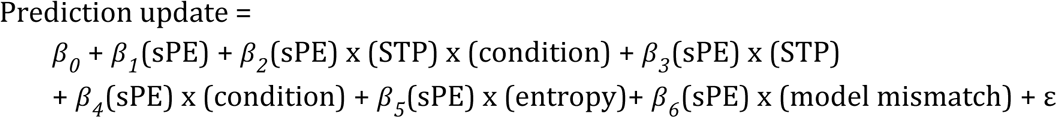

Where STP stands for state transition probability and condition is a binary dummy variable with changepoint assigned as 1 and oddball assigned as -1. *β_1_* is a term that captures the average rate of learning. *β_2_* is an interaction term between STP, sPE, and condition that captures the bidirectional learning effect predicted by our model: increased learning for state transitions in the changepoint condition and decreased learning for state transitions in the oddball condition. *β_3_* measures how STP alone changes the slope of average learning rate. The error term, ε, is assumed to be independent and von Mises distributed to account for the nature of the circular color wheel in our task.

The model also included nuisance variables that captured variance unrelated to our primary hypotheses. *β_4_* captures how the slope of average learning rate adjusts based on task condition. Entropy captures trial-to-trial uncertainty. *β_5_* describes how entropy affects the slope of average learning rate. Model mismatch captures trials where subject predictions diverge from model predictions. To calculate this term, we fit the differences between model and subject prediction for each trial under the same task structure with a mixture of gaussian and uniform distributions with mixture proportion and precision as free parameters. Model mismatch is the likelihood for each trial that subject report was uniform.

To examine perception, we measured trial-to-trial color reproduction error as: ground truth color(t) – color reproduction(t). Perceptual bias was measured at the individual subject level as the slope of the relationship between reproduction error and PE. PE, in this case, was defined as the objective prediction error (oPE), which is the difference between stimulus color prediction and the ground truth of the same stimulus color on the same trial. To understand how perceptual bias was affected by state transitions, we performed the following regression on both the model and subject behavior data:

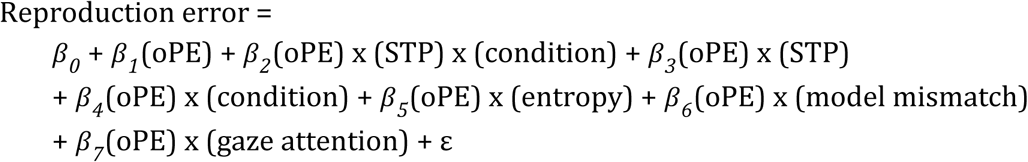

Where *β*_1_captures the overall degree of perceptual bias, and *β*_3_ captures the degree to which such biases are suppressed at likely state transitions. Note this regression is similar to the regression for learning, with the additional *β_7_* term and the dependent variable changed to reproduction error and prediction error term changed to oPE. The nuisance variable (*β_7_*) captures the degree to which average horizontal gaze position during stimulus onset (0-200ms) affects the relative bias in perceptual reports related to the right and left target colors. The gaze attention variable is zero for both targets when fixated, negative if gaze is toward the target and positive if gaze is away from the target. The coefficient (*β_7_*) estimates the contribution of the interaction between the relative gaze attention and prediction errors on perceptual errors, allowing us to capture variability in bias that might be related to differences in visual attention.

### Pupil & EEG Data Analysis

Both pupil area and EEG signals were regressed onto an explanatory matrix that contained the following:

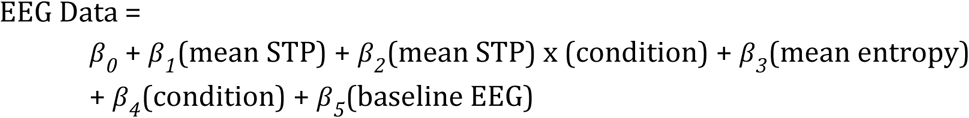

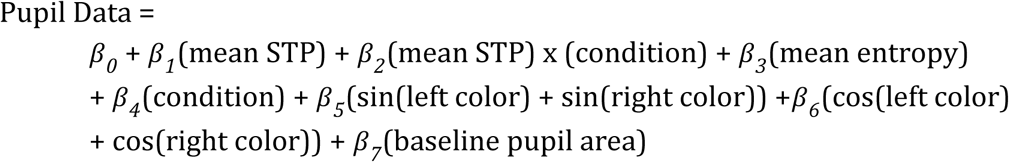

To further minimize the effects of the presented color on pupil dilation, we also included the sum of the sines and cosines of stimulus colors as nuisance regressors in the pupil data regression. Regression coefficients were combined across subjects to compute t-statistics at each time point (and channel in the case of EEG). T-statistics were thresholded with a cluster-forming threshold corresponding to p=0.01 for EEG data and p=0.025 for pupil data, and temporal or spatiotemporal clusters were submitted to permutation testing to identify clusters with mass (summed T-statistic within cluster) that exceeded 97.5% of values in the permuted null distribution.

Trial-to-trial pupil and EEG signal strength was quantified within windows of significant association with STP as the dot product of baseline-regressed individual trial responses and the thresholded t-statistic map over time (for pupil analysis) or time and channels (for EEG analysis). Trial-to-trial measures of pupil and EEG response were incorporated into two regression models to explain learning rate and bias in the changepoint and oddball blocks. These regressions were run for each subject separately and coefficients for all participants were compared against zero using a student’s t-test to determine significance of relationships between learning/bias and arousal. In addition to this statistical testing, we also qualitatively tested our latent state predictions. To visualize our results, we sorted pupil dilation and P300 data into 5 quantiles and computed and plotted the perceptual bias and learning rates as described above separately in each quantile.

We also examined pupil size during the prediction phase of the task in two different time windows in order to test whether oddballs promote higher levels of arousal on the subsequent trial than changepoints. Pupil size in a time window beginning one second before the end of the color estimation phase and ending five seconds later was regressed onto an explanatory matrix. This explanatory matrix was as follows:

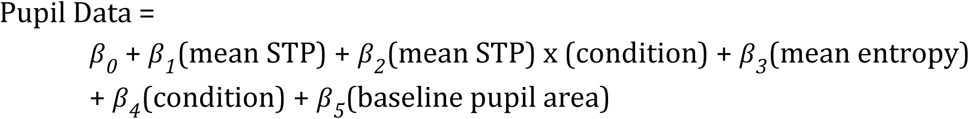

Note that the color terms were removed from this regression because the time window we were investigating did not include stimulus presentation. Significance was assessed by computing t-statistics across subjects and controlling for multiple comparisons using permutation testing as described above.

We also assessed pupil size during a six second window from two seconds before each prediction was made during a trial to four seconds after the prediction was made. Since participants’ eye data was lower quality during this more demanding period of the task and participants were often squinting, we removed any trials where a participant’s pupil size was greater than six standard deviations smaller than the participant’s mean pupil size. Pupil size during this time window was regressed onto an explanatory matrix that contained the following:

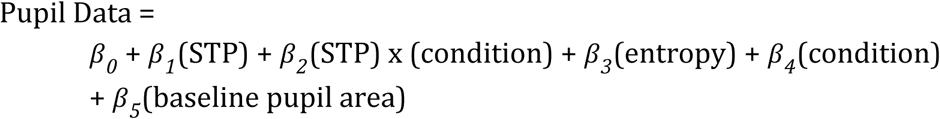

The STP and entropy terms for this regression were not averaged because they were calculated at the time a prediction was made. Thus, if the left stimulus was prompted, the STP term would be the STP value for the left stimulus only rather than the mean STP across both stimuli. The baseline time window was from 1500 to 500 ms before the prediction was made. To determine whether coefficients in this regression were statistically significant across time, we performed a permutation test identical to that done on the previous pupil regression coefficients. To assess whether the STP * condition coefficient was related to behavior, we correlated the eye regression coefficients with the regression from the behavioral regression on prediction update. To quantify each subject’s degree of dilation when predicting after oddballs, we calculated each subject’s average STP * condition regression coefficient during the period that was deemed to be significantly negative by permutation testing. We then performed a Pearson correlation on each subject’s average STP * condition coefficient with the sPE * STP * condition coefficient from the previous update regression.

Since the timing of this event was made somewhat unclear by our baselining process, we also performed the same analysis on the derivative of the pupil area during the same time window (2 seconds before prediction to 4 seconds after prediction). We convolved the pupil signal across a window of 150 ms, took the derivative of the convolved signal, and ran the same running regression across those data. Significance was assessed by performing a t-test across each participant’s STP x condition coefficient, then performing a permutation test to assess temporal clusters of significant p-values.

### Joint Pupil-EEG Analysis

For analyses using a single value for arousal, pupil area and EEG data were combined. The individual trial scores for both the EEG and pupil data were (in the case of EEG) summed, then normalized to create one value each for pupil and EEG that were weighted equally. These values were then summed to create a single value corresponding to arousal during the trial. Any trials that were rejected before in the EEG data or had a blink percentage higher than 50% were removed from the analysis. We then performed the same regression on the joint pupil-EEG arousal signal that we did on the separate EEG and pupil signals to quantify the relationship between arousal and learning or bias. To visualize the joint arousal data, we sorted each block of subject data into 5 quantiles and plotted the mean learning rate or bias across subjects for each quantile, similarly to how we visualized the separated pupil and EEG data.

To quantify the relationship between learning/arousal slopes and bias/arousal slopes across both blocks, we calculated the bias/arousal slope combined across blocks and plotted it against the learning/arousal slopes for each block separately that had been previously calculated from the joint pupil-EEG data. Each bias/arousal value had two coordinates associated with it, as it was plotted against the values calculated for the subject’s changepoint and oddball learning/arousal slope separately. We then performed a regression on the learning/arousal values as follows:

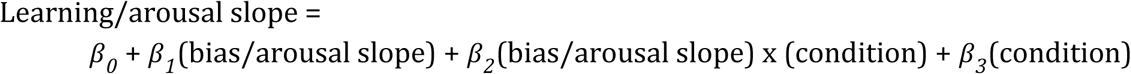

To determine a relationship between bias/arousal slope and learning/arousal slope, we looked for a significant value in *β_2_*, a negative relationship would show that subjects with stronger learning/arousal slopes tended to have stronger bias/arousal slopes in both blocks.

We also wanted to investigate whether the relationships we found in the earlier investigations were robust even if STP was removed. To do this we regressed mean trial STP from the trial average learning rates, trial average bias, and trial-by-trial EEG and pupil signals. Then, we performed the pupil and EEG data combination method we used described above and repeated our analyses above to calculate learning/arousal and bias/arousal relationships in both blocks. We then performed the regression of STP-regressed bias/arousal slope against STP-regressed learning/arousal slope as described above to determine whether the relationship between bias slope and learning slope survived after regressing out STP.

## Supplemental Figures

**Figure S1:**
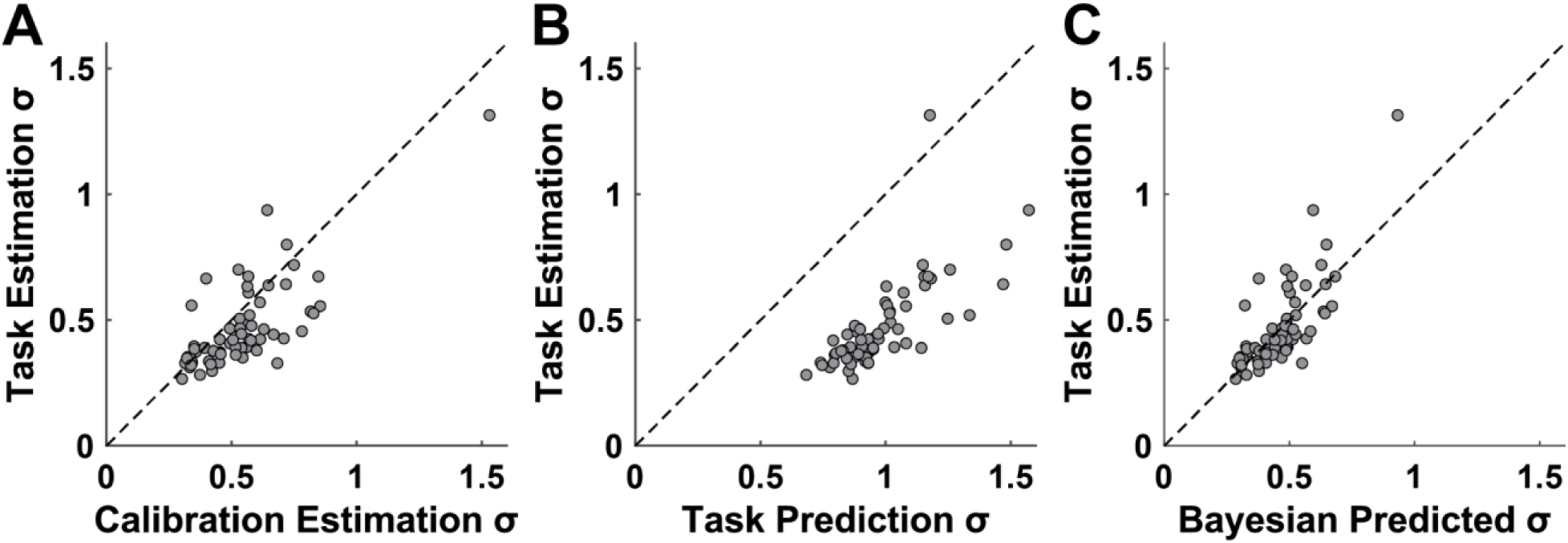
Subjects in structured blocks are able to use perceptual bias to improve perception. **(A)** The standard deviation of estimation error in the practice phase is significantly higher than in the prediction phase (t = 4.74; dof = 62; p = 6.45*10^−6^). **(B)** Prediction error standard deviation is much larger than estimation error standard deviation (t = 24.3; dof = 62; p = 2.26*10^−33^) **(C)** Participant estimation error distribution in the prediction phase is not statistically different from the prediction error distribution multiplied by the estimation error distribution in the practice phase (t = -0.95; dof = 62; p = 0.348)

**Figure S2:**
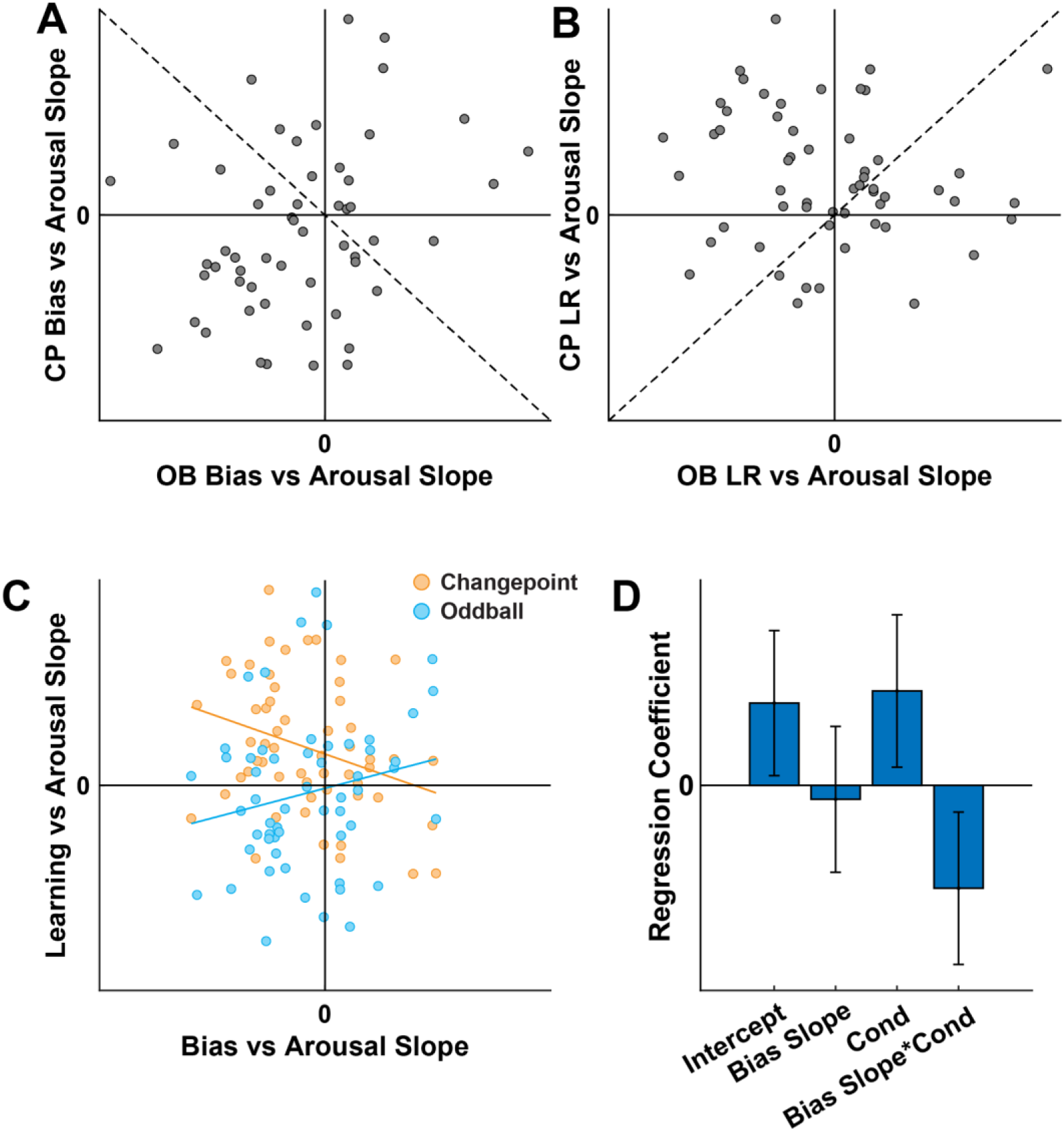
Behavioral relationships to state transition signals are partially maintained even when STP is removed from learning rate, bias, and arousal signals. **(A)** After regressing out STP, changepoint bias/arousal slope is not significant (t = -1.41; dof = 55; p = 0.1632), OB bias/arousal slope is negative on average (t = -2.49; dof = 55; p = 0.0158), and the sum of CP and OB bias/arousal slope is negative (t = -2.38; dof = 55; p = 0.0207). **(B)** After regressing out STP, changepoint learning/arousal slope is positive on average (t = 4.23; dof = 55; p = 8.85*10^−5^), OB learning/arousal slope is not different from zero (t = -0.72; dof = 55; p = 0.4745), and the difference between CP and OB learning/arousal slopes is positive (t = 3.11; dof = 55; p = 0.0030). **(C)** Individual differences plot on STP-regressed residuals. Blue points indicate subjects’ slopes in the oddball lock, orange points indicate slopes in the changepoint block. While learning vs arousal slopes were individually calculated within each block, bias vs arousal slopes were calculated across both blocks. **(D)** Regression results against learning slopes. The negative effect of bias slope * condition suggests the relationship between learning/arousal slope and bias/arousal slope is partially maintained even when STP is removed (Bias slope * condition t = -2.67; dof = 108; p = .0087).

**Figure S3:**
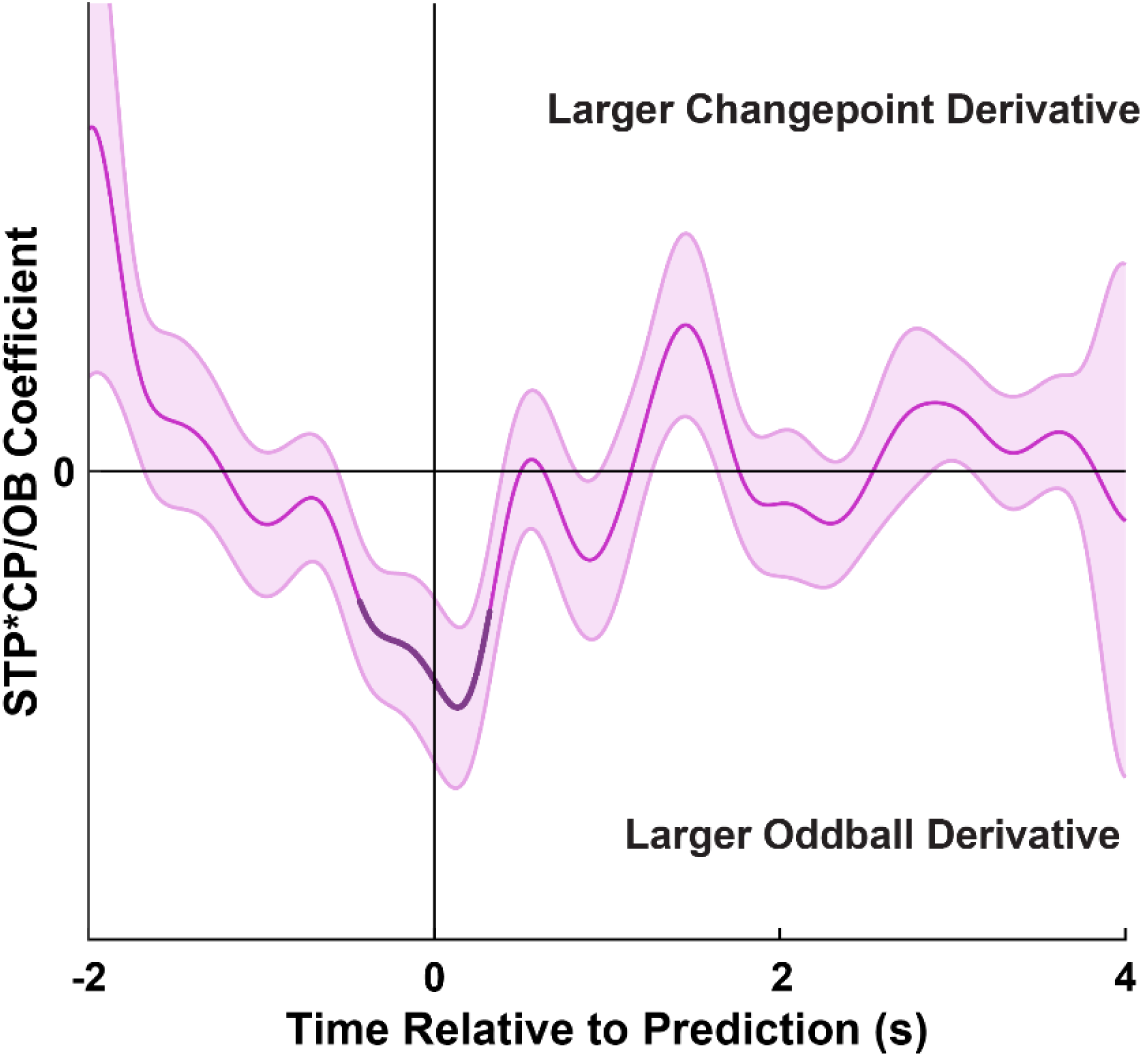
Derivative analysis of pupil dilation at the time of prediction shows dilation starts before predictions are selected. Mean/SEM STP*condition coefficients from a running linear regression of pupil size derivative locked to the prediction phase of the trial. Dark points reflect significant clusters after permutation testing for multiple comparisons correction (STP*CP/OB cluster forming threshold = 0.025, Cluster mass = 1940, p = 0.0102). The significant coefficient before the time of prediction indicated that a state transition signal from LC would have to arise before the prediction as well.

**Figure S4:**
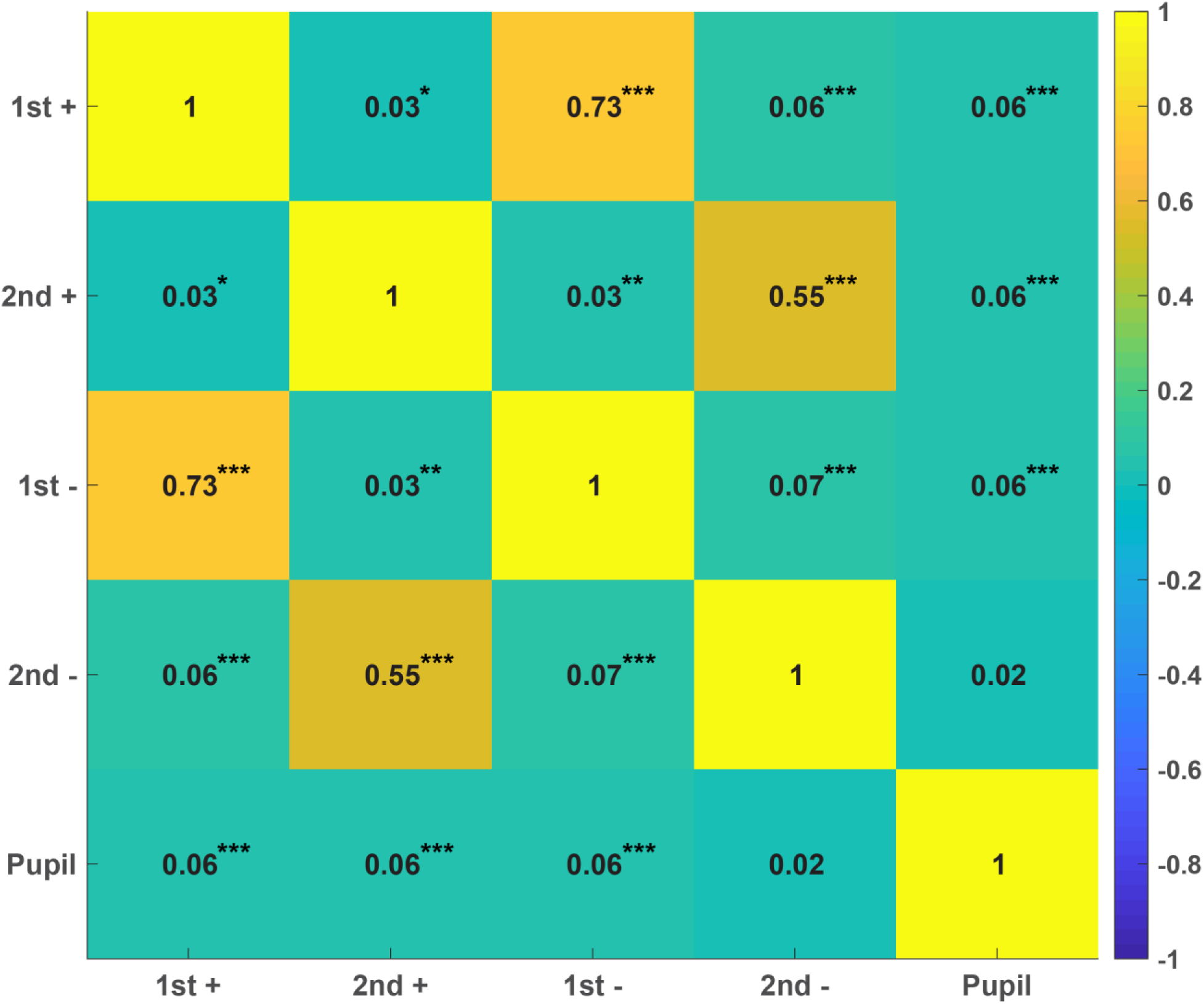
Correlation between individual EEG and pupil signals: Average correlation coefficients across subjects between different components of the state transition signal (1^st^ positive P3a-like signal, second frontal positive, 1^st^ parietal negative, second parietal negative, pupil dilation), values marked * indicate p<0.05, ** indicate p<0.01, *** indicate p<0.001.

**Figure S5:**
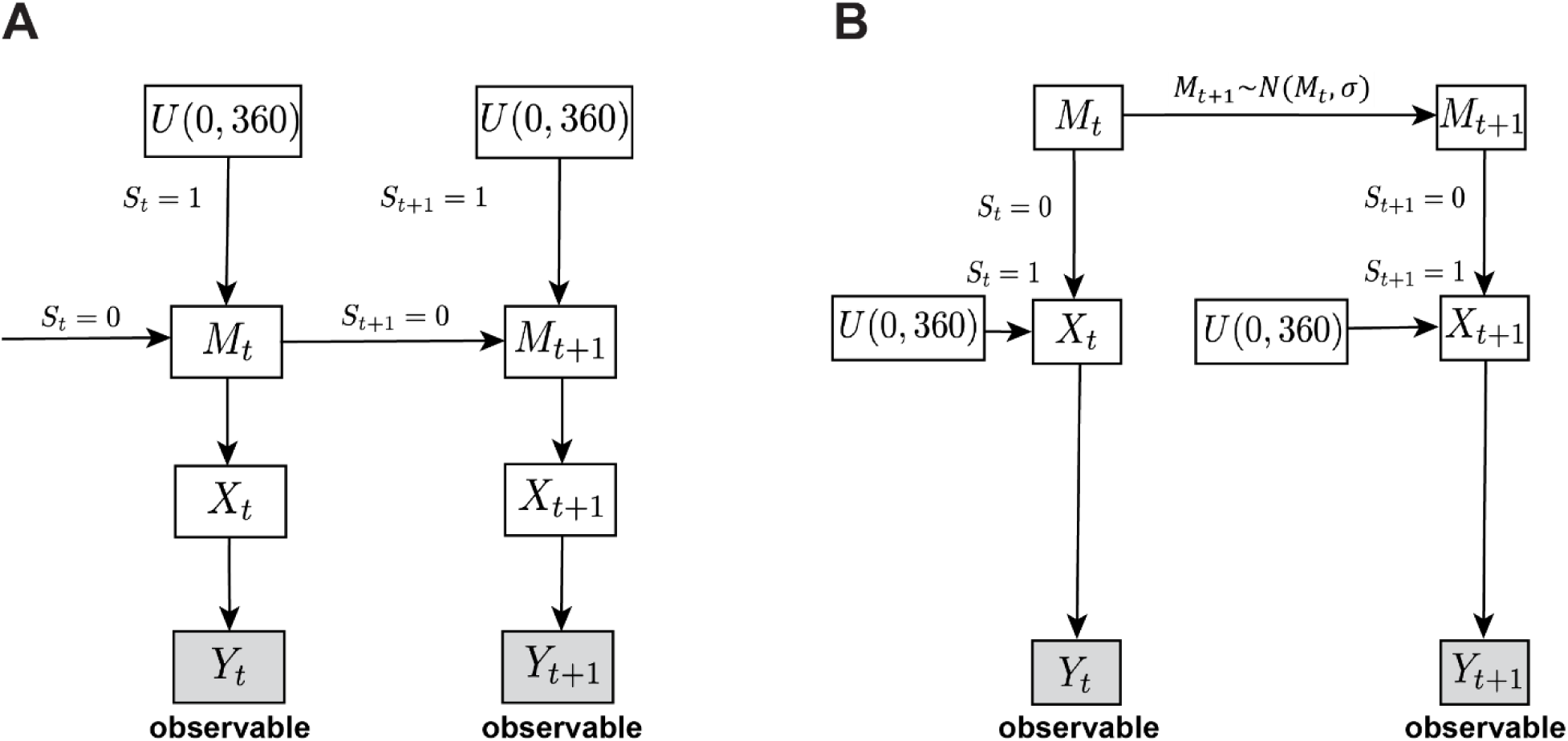
Task generative model creates changepoint and oddball structures. **(A)** Generative model for the changepoint block. Each trial’s presented color, *X*, is generated about the underlying mean, *M*. On non-changepoint trials, *M* remains constant from previous trials. On changepoint trials, *M* is sampled randomly from a uniform distribution about the color space. **(B)** Generative model for the oddball block. On normal trials, presented color, *X*, is generated about an underlying mean, *M*. On oddball trials, *X* is selected from a uniform distribution about the color space. Between each trial, *M* drifts with variance, *σ*.

